# An Off-the-Shelf Bioadhesive Patch for Sutureless Repair of Gastrointestinal Defects

**DOI:** 10.1101/2021.03.12.435203

**Authors:** Jingjing Wu, Hyunwoo Yuk, Tiffany L. Sarrafian, Chuanfei Guo, Leigh G. Griffiths, Christoph S. Nabzdyk, Xuanhe Zhao

**Affiliations:** Department of Mechanical Engineering, Massachusetts Institute of Technology, Cambridge, MA, USA; Department of Thoracic Surgery, Mayo Clinic, Rochester, MN, USA; Department of Materials Science and Engineering, Southern University of Science and Technology, Shenzhen, China; Department of Cardiovascular Diseases, Mayo Clinic, Rochester, MN, USA; Department of Anesthesiology and Perioperative Medicine, Mayo Clinic, Rochester, MN, USA; Department of Civil and Environmental Engineering, Massachusetts Institute of Technology, Cambridge, MA, USA

## Abstract

Surgical sealing and repair of injured and resected gastrointestinal (GI) organs are critical requirements for successful treatment and tissue healing. Despite being the standard of care, hand-sewn closure of GI defects using sutures faces various limitations and challenges. The process remains technically complicated and time-consuming. The needle-piercing and pointwise closure also inflict tissue damage and stress concentration, raising the risk of local failure and subsequent anastomotic leaks. To address these limitations and challenges, we introduce an off-the-shelf bioadhesive GI patch capable of atraumatic, rapid, robust, and sutureless repair of GI defects. The GI patch synergistically integrates a non-adhesive top layer and a dry bioadhesive bottom layer, resulting in a thin, flexible, transparent, and ready to use dressing with tissue-matching mechanical properties. Rapid, robust, and sutureless sealing capability of the GI patch is systematically characterized based on various standard tests in *ex vivo* porcine GI organ models. *In vitro* and *in vivo* rat models are utilized to validate biocompatibility and biodegradability of the GI patch including comprehensive cytotoxicity, histopathology, immunofluorescence, and blood analyses. To validate the GI patch’s efficacy in a clinically relevant setting, we demonstrate successful sutureless *in vivo* sealing and healing of GI defects; namely in rat stomach and colon, and porcine colon injury models. The proposed GI patch not only provides a promising alternative to suture for repair of GI defects but also offers potential clinical opportunities in the treatment and repair of other organs.

**One Sentence Summary:** An off-the-shelf bioadhesive patch is introduced for facile sutureless repair of gastrointestinal defects, addressing various limitations of conventional suture-based treatments.

## INTRODUCTION

Failure of surgical repair of gastrointestinal (GI) defects, can lead to anastomotic leaks, one of the most feared and life-threatening complications following GI surgeries, resulting in an over 30 % increase in mortality (*1, 2*). Surgical sealing and repair of GI defects is commonly achieved by using sutures and perform hand-sewn closure of GI tissues. However, despite being the standard of care, GI organ sealing through sutures remains associated with a high rate of anastomotic leaks (i.e., up to 20 % in high-risk patients), resulting in serious complications including infection, sepsis, and even death (*1–5*). While failure in surgically repaired GI tissues can be due to various factors (*1, 6*), one major culprit for the high failure rate has been the inherent disadvantages of suture-based tissue sealing including (i) complicated technical processes that requires high surgical skill, (ii) tissue damage due to needle-piercing nature, and (iii) point-wise closure leading to stress-concentration around the sutured points (Fig. 1A) (*2, 3, 5, 7, 8*). As an alternative to sutures, surgical staplers have been increasingly adopted for GI surgeries. However, surgical staplers also face similar limitations such as tissue damage, pointwise closure and stress concentration, and they do not significantly reduce the rate of leaks compared to sutures (*9*). Hence, surgical repair of GI defects to provide mechanical sealing and favorable healing still remains an ongoing challenge in the field, highlighting the critical importance of developing new treatments and solutions.

**Fig. 1.**
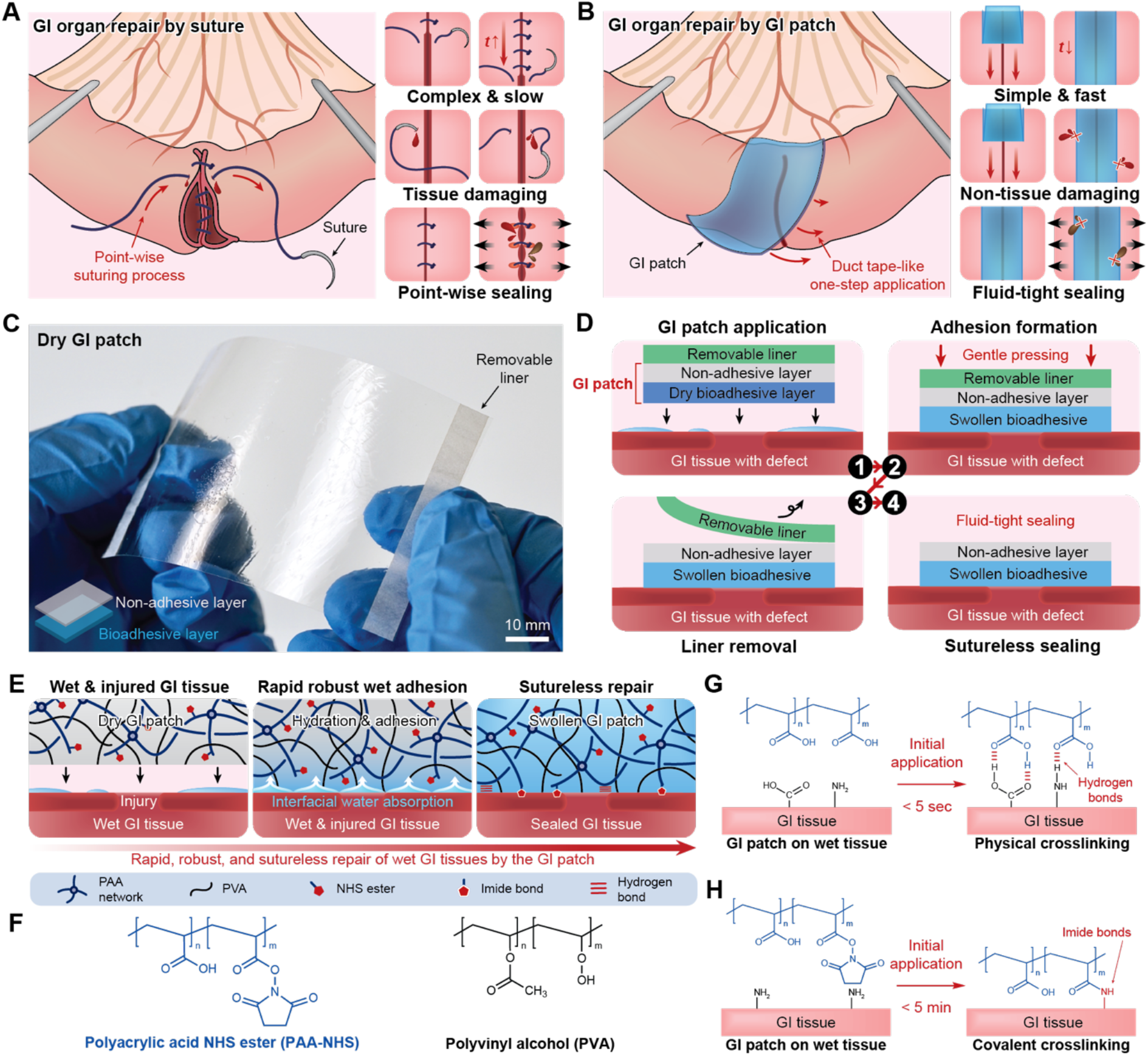
Design and mechanism of sutureless repair by the GI patch. (**A**) Schematic illustrations for repair of GI defects by sutures. (**B**) Schematic illustrations for sutureless repair of GI defects by the GI patch. (**C**) Image of the dry GI patch and a schematic illustration (bottom-left) for its structure consisting of the non-adhesive top layer and the bioadhesive bottom layer. (**D**) Schematic illustrations for the components of the GI patch and for the step-wise processes of sutureless repair of GI defects by the GI patch. (**E**) Schematic illustrations for mechanism of sutureless repair of GI defects by the GI patch based on the dry-crosslinking process. (**F**) Chemical composition of the GI patch based on poly(acrylic acid)-NHS ester (PAA-NHS) and poly(vinyl alcohol) (PVA). (**G** and **H**) Schematic illustrations for rapid wet adhesion of the GI patch to the GI tissue surface by physical crosslinking based on hydrogen bonds (G) and covalent crosslinking based on imide bonds (H). Scale bars, 10 mm (C).

Tissue adhesives and sealants have recently emerged as an alternative or adjunct to sutures or staples in various clinical indications, owing to their potential advantages (*7, 8, 10–17*). However, existing tissue adhesives and sealants are mostly deployed in the form of viscous liquids, and they usually require a diffusion-based interpenetration into tissues (*11, 13, 14*) and/or solidification by chemical reaction or external stimuli such as ultraviolet (UV) light (*13–15* to form tissue sealing. These features of existing tissue adhesives and sealants result in several limitations including slow and/or weak tissue sealing, the need for external devices (e.g., UV source or fluidic mixer), and/or complicated preparations and applications (e.g., thawing, mixing or light irradiation) (*13, 14, 18*), which render them far from ideal for facile and robust repair of GI defects. More recently, to overcome the limitations of existing technologies, several bioadhesives have been developed to provide rapid and/or tough adhesion to wet tissues (*19–23*). However, the materials used and their underlying sealing mechanisms have not sufficiently optimized for such challenging clinical applications as the repair of GI defects.

In this study, we introduce an off-the-shelf bioadhesive patch platform, named GI patch, to offer a promising therapeutic solution for the treatment of GI defects (Fig. 1B). Inspired by the convenience and effectiveness of duct-tape in non-medical applications, the GI patch offers facile, atraumatic, fluid-tight, robust, and sutureless sealing of GI defects, while addressing key limitations of sutures and commercially-available tissue adhesives and sealants (Fig. 1B). The GI tissue-matching mechanical properties and superior adhesion performance of the GI patch are systematically characterized by various standard tests based on *ex vivo* porcine models. The biocompatibility and biodegradability of the GI patch is thoroughly validated through *in vitro* cytotoxicity evaluation in human intestinal epithelial cells, and *in vivo* histopathology, immunofluorescence, and blood analyses in rat models. We further validate the *in vivo* efficacy of sutureless repair of GI defects by the GI patch in rat stomach and colon injury models and a porcine colon injury model.

## RESULTS

### Design and mechanisms of the GI patch

The GI patch takes the form of a thin, flexible, and transparent dressing consisting of a nonadhesive top layer and a dry bioadhesive bottom layer (Fig. 1C and fig. S1). A removable liner layer can be further assembled on top of the non-adhesive layer if the liner may improve the handling of the GI patch in certain applications (Fig. 1C and D). The non-adhesive top layer consists of hydrophilic polyurethane to provide a hydrophilic and non-adhesive interface to the surrounding tissues while providing tissue-matching and robust mechanical properties to the GI patch. The dry bioadhesive bottom layer consists of interpenetrating networks between the covalently-crosslinked poly(acrylic acid) NHS ester (PAA-NHS) for bioadhesiveness and the physically-crosslinked poly(vinyl alcohol) (PVA) for mechanical reinforcement (Fig. 1E and F). The bioadhesive can form fast and robust adhesion on wet GI tissues based on a dry-crosslinking mechanism (*20, 21, 24*). In brief, the hydrophilic and hygroscopic PAA-NHS and PVA in the dry bioadhesive layer allows the absorption and cleaning of the interfacial water on wet GI tissues upon contact (*20, 24*) (Fig. 1E). Subsequently, the carboxylic acid groups and NHS ester groups in the PAA-NHS network facilitate rapid and robust adhesion to the GI tissue surface based on physical crosslinking via hydrogen bonds (Fig. 1G) and covalent crosslinking via imide bonds (Fig. 1H), respectively (*20, 21, 25*). The off-the-shelf flexible dressing form factor and the facile adhesion to wet GI tissues without the need of other devices or stimuli (e.g., UV) synergistically endow the GI patch with unique ready-to-use and preparation-free features, similar to those features of duct-tapes.

After adhering to GI tissues and sealing GI defects, the GI patch is further hydrated and swells in a wet physiological environment (fig. S2) into a soft (Young’s modulus of 135 kPa), stretchable (over 5 times of the original length, fig. S3A), and robust (fracture toughness of 758 J m^-2^, fig. S4) hydrogel. Notably, the swelling-driven lateral dimensional changes (i.e., length and width directions) of the GI patch have been eliminated (fig. S2) by introducing a pre-strain to the dry bioadhesive layer equivalent to its swelling ratio during the GI patch preparation (fig. S1). This unique characteristic of the GI patch prevent separation of approximated wound edges and subsequent delayed healing, which is a common problem in various swellable tissue adhesives and sealants (*8*). To minimize the GI patch’s mechanical mismatch with GI tissues, the Young’s modulus of the GI patch is optimized to match that of GI organs (fig. S3D), based on the experimentally measured toe Young’s moduli of *ex vivo* porcine colon and stomach (fig. S3B and C). Furthermore, the GI patch maintains consistent mechanical properties in terms of Young’s modulus, ultimate tensile stretch, and tensile strength up to one month in physiological environments (fig. S5), potentially offering mechanical stability and integrity during the critical stages of GI tissue defect healing.

### Adhesion performance

To quantitatively evaluate the GI patch’s capability to form rapid and robust adhesion to GI tissues, we characterize interfacial toughness, shear strength, tensile strength, and burst strength of the GI patch in *ex vivo* porcine colon and stomach tissues (Fig. 2 and fig. S6). The GI patch can provide facile and robust adhesive sealing of GI tissues upon contact for 5 s with high interfacial toughness (over 350 J m^-2^ for colon; over 500 J m^-2^ for stomach), shear strength (over 65 kPa for colon; over 80 kPa for stomach), and tensile strength (over 60 kPa for colon; over 65 kPa for stomach) (Fig. 2A-F). Notably, the GI patch exhibits superior adhesion performance in comparison to various commercially-available tissue adhesives and sealants including cyanoacrylate glue (Histoacryl^®^), poly(ethylene glycol) (PEG)-based sealant (Coseal), and fibrin glue (Tisseel) for both porcine colon (Fig. 2A-C) and stomach (Fig. 2D-F).

**Fig. 2.**
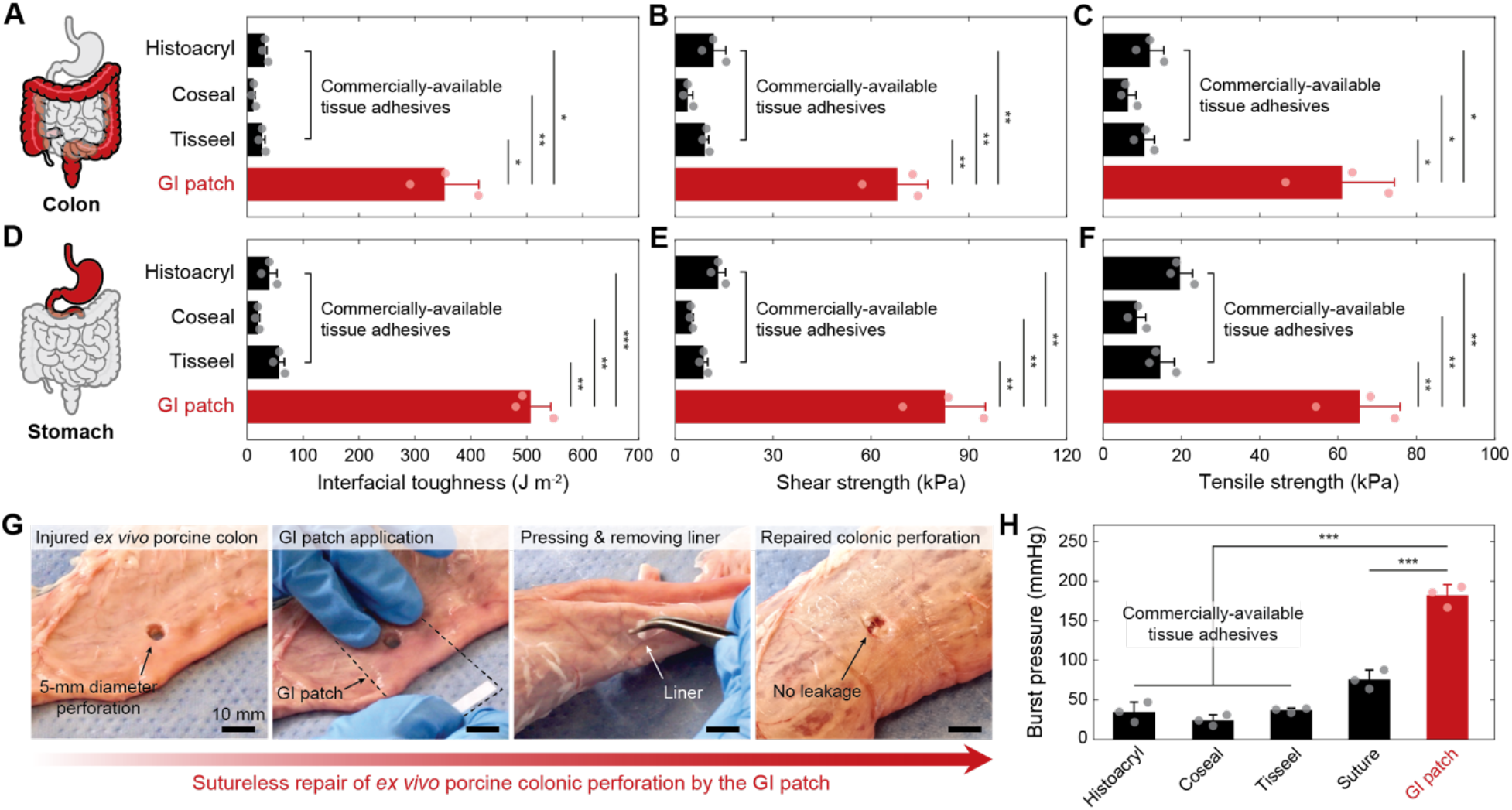
Adhesion performance of the GI patch. (**A-C**) Adhesion performance of the GI patch and the commercially-available tissue adhesives for interfacial toughness (A), shear strength (B), and tensile strength (C) on *ex vivo* porcine colon. (**D-F**) Adhesion performance of the GI patch and the commercially-available tissue adhesives for interfacial toughness (D), shear strength (E), and tensile strength (F) on *ex vivo* porcine stomach. (**G**) Snapshots of rapid and robust sutureless repair of 5-mm diameter defects in an *ex vivo* porcine colon by the GI patch. (**H**) Burst pressure of an *ex vivo* porcine colon with a 5-mm diameter defect sealed by the GI patch, the commercially-available tissue adhesives, and sutures. Values in A-F and H represent the means ± SD (*n* = 3). *P* values are determined by the Student’s *t* test for the comparisons between two groups and by one-way ANOVA followed by the Tukey’s multiple comparison test for the comparison between multiple groups; * *p* ≤ 0.05; \*\**p* ≤ 0.01; \*\*\**p* ≤ 0.001. Scale bars, 10 mm (G).

We further evaluate sutureless fluid-tight sealing of GI defects by the GI patch based on *ex vivo* porcine colon and stomach models (Fig. 2G). Enabled by rapid and robust adhesion to GI tissues, the GI patch can readily form fluid-tight sealing of 5-mm diameter defects in porcine colon (movie S1) and stomach (movie S2) in less than 10 s after application, demonstrating the sutureless sealing capability of the GI patch. Moreover, the seal formed by the GI patch exhibits high burst pressure of over 180 mmHg, significantly outperforming the commercially-available tissue adhesives and sealants (Histoacryl^®^, Coseal, Tisseel) as well as sutures (Fig. 2H).

### Biocompatibility and biodegradability

*In vitro* LIVE/DEAD staining of human intestinal epithelial cells (Caco-2) cultured in GI patch-incubated media for 24 h shows comparable cell viability to the control media group (*p* = 0.12), whereas exposure to cell media incubated with the commercially-available tissue adhesive groups (Coseal and Histoacryl^®^) significantly lowers cell viability compared to the control media group (Fig. 3A). *In vivo* biocompatibility of the GI patch and the commercially-available tissue adhesives (Coseal and Histoacryl^®^) are assessed based on the implantation onto intact rat colon and stomach for 4 weeks (fig. S7), followed by histopathology, immunofluorescence, and blood analyses (Fig. 3B-J). The histological evaluation by a blinded pathologist indicates that the GI patch induces minimal to no inflammation to the underlying and surrounding GI tissues (Fig. 3F and G), comparable to that of the Coseal group (Fig. 3B and C). In contrast, cyanoacrylate glue (Histoacryl^®^) exhibits lower *in vivo* biocompatibility than the GI patch and Coseal groups with mild inflammation and fibrosis (Fig. 3D and E), agreeing with the previous reports (*22*). Notably, the Histoacryl^®^ group shows marked fibrotic adhesion to the surrounding tissues (fig. S7B and E) while such macroscopic fibrotic adhesion is not observed in the GI patch and Coseal groups.

**Fig. 3.**
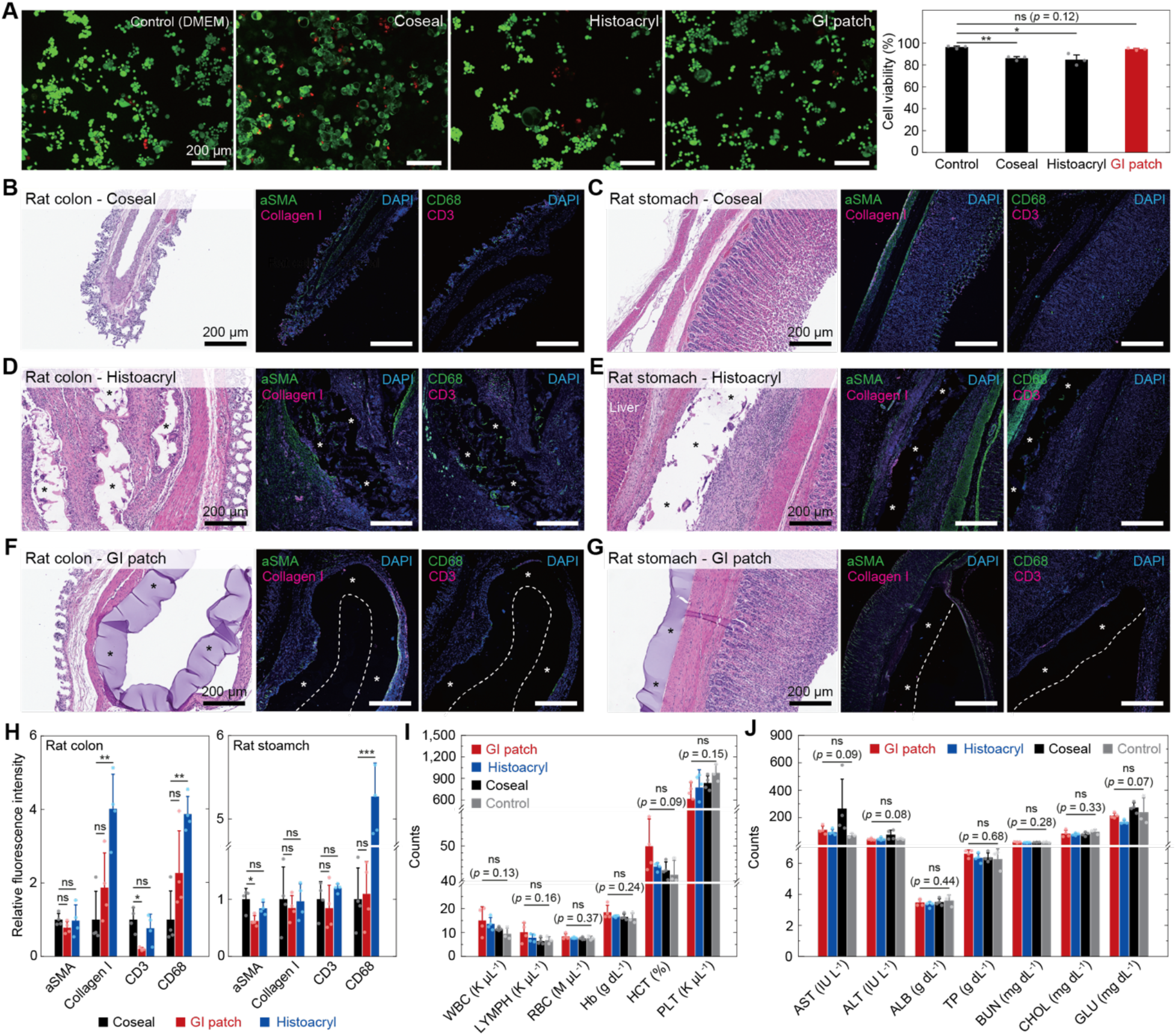
*In vitro* and *in vivo* biocompatibility of the GI patch. (**A**) Representative LIVE/DEAD assay images (left) and the cell viability (right) of human intestinal epithelial cells (Caco-2) for control (DMEM), Coseal, Histoacryl, and the GI patch after 24-h culture. DMEM, Dulbecco’s Modified Eagle Medium. (**B-G**) Representative histological images stained with hematoxylin and eosin (HE) (left) and immunofluorescence images (right) for Coseal implanted to rat colon (B) and rat stomach (C); Histoacryl implanted to rat colon (D) and rat stomach (E); the GI patch implanted to rat colon (F) and rat stomach (G) for 4 weeks. In histological images, * represents the implanted Histoacryl (D and E) and GI patch (F and G). In immunofluorescence images, blue fluorescence corresponds to cell nuclei stained with 4′,6-diamidino-2-phenylindole (DAPI); green fluorescence corresponds to the expression on fibroblast (αSMA) and macrophages (CD68); red fluorescence corresponds to the expression on collagen (Collagen I) and T-cell (CD3); * represents the implanted Histoacryl (D and E) and GI patch (F and G); dotted line represents the edge of the implanted GI patch. (**H**) Normalized fluorescence intensity from the immunofluorescence images for αSMA, Collagen I, CD3, and CD68 after 4 weeks implantation of Coseal, Histoacryl, and the GI patch to rat colon (left) and stomach (right). (**I**) Complete blood count (CBC) of control (healthy animal without surgery) and after 4 weeks implantation of Coseal, Histoacryl, and the GI patch to rat colon and stomach. WBC, white blood cell; LYMPH, lymphocyte; RBC, red blood cell; Hb, hemoglobin; HCT, hematocrit; PLT, platelet. (**J**) Blood chemistry of control (healthy animal without surgery) and after 4 weeks implantation of Coseal, Histoacryl, and the GI patch to rat colon and stomach. AST, aspartate transaminase; ALT, alanine transaminase; ALB, albumin; TP, total protein; BUN, blood urea nitrogen; CHOL, cholesterol; GLU, glucose. Values in A and H-J represent the means ± SD (*n* = 3 in A; *n* = 4 in H-J). *P* values are determined by the Student’s *t* test for the comparisons between two groups and by one-way ANOVA followed by the Tukey’s multiple comparison test for the comparison between multiple groups; ns, not significant; * *p* ≤ 0.05; ** *p* ≤ 0.01; *** *p* ≤ 0.001. Scale bars, 200 μm (A-G).

The immunofluorescent staining (Fig. 3B-G) and the normalized immunofluorescence intensity analysis (Fig. 3H) of fibroblasts (αSMA), Collagen I, macrophages (CD68), and T-cells (CD3) further confirm that the GI patch induces inflammatory and foreign body responses comparable to the Coseal group and significantly less than the Histoacryl^®^ group. The blood analysis including complete blood counts (CBC) and comprehensive blood chemistry panels shows that all three groups (the GI patch, Coseal, Histoacryl^®^) are comparable to the healthy rat control group without notable systemic toxicity for the study period of 4 weeks (Fig. 3I and J).

We further investigate the *in vivo* biodegradability of the GI patch in a rat subcutaneous implantation model for up to 12 weeks (fig. S8). The histological assessment at 2, 4, 8, and 12 weeks after the implantation indicates that the GI patch maintains its film-like configuration until 4 weeks while exhibiting notable reduction in the thickness at 8 weeks (fig. S8). At the longer time points such as 12 weeks, the GI patch shows fragmentation and degradation, supporting the biodegradability of the GI patch (fig. S8).

### Sutureless repair and healing of GI defects in rat models

To validate the *in vivo* efficacy of sutureless repair of GI defects by the GI patch, we evaluate rapid, robust, and fluid-tight sealing and healing of rat colon and stomach defects by the GI patch in comparison with sutures as a standard care control (Fig. 4A and F). Notably, the GI patch can be prepared into a ready-to-use package both with and without a removable liner as a backing substrate (fig. S1B), and the GI patch with a removable liner is used for the sutureless repair of GI defects in the rat models to facilitate easier handling of the thin GI patch for small animals. We demonstrate that the GI patch can readily establish atraumatic and fluid-tight sealing of 10-mm incisional defects in rat colon (Fig. 4B and movie S3) and stomach (Fig. 4G and movie S4) in less than 10 s without further preparation steps or the need of additional devices (e.g., mixer, UV light). In contrast, the point-wise tissue closure by sutures takes much longer time (over 2 minutes) and causes puncture-driven tissue damages (fig. S9). In comparison, the GI patch offers simple, consistent, and fully conformal adhesive sealing of the GI defects for all rats in the study. After 4 weeks post-repair, both groups (the GI patch, sutures) show healing of the defects without macroscopic sign of GI leaks. Notably, the GI patch maintains robust adhesion to the underlying GI tissues in all rats in the study at the moment of sacrifice (i.e., 4 weeks) (Fig. 4C and H).

**Fig. 4.**
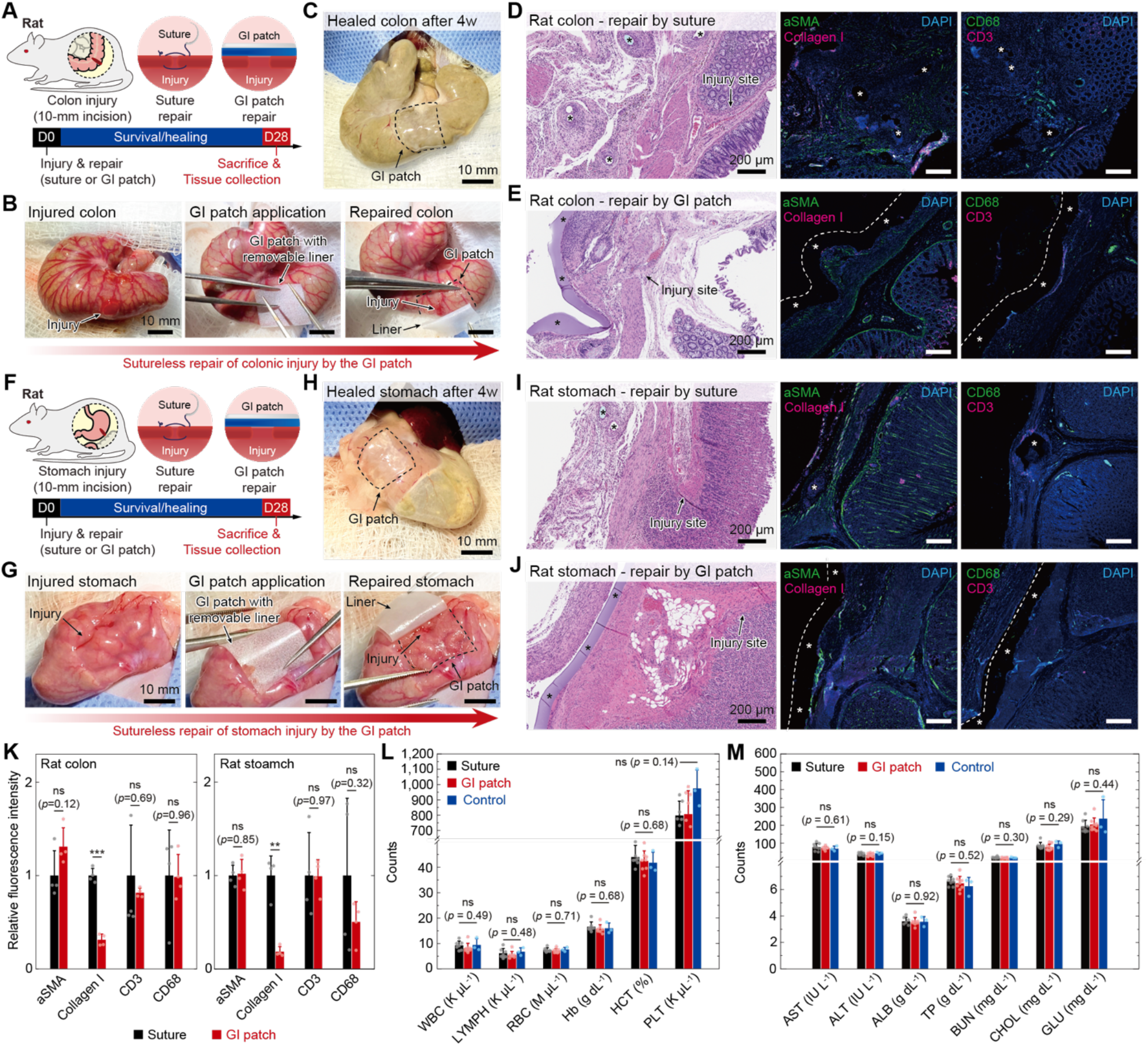
Sutureless repair of GI defects in rat model. (**A** and **B**) Schematic illustrations (A) and experimental images (B) for *in vivo* defect-repair studies of rat colon by sutures and the GI patch. (**C**) Rat colon 4 weeks after sutureless repair by the GI patch. (**D** and **E**) Representative histological images stained with hematoxylin and eosin (HE) (left) and immunofluorescence images (right) for rat colon defect repaired by sutures (D) and the GI patch (E) after 4 weeks. (**F** and **G**) Schematic illustrations (F) and experimental images (G) for *in vivo* defect-repair studies of rat stomach by sutures and the GI patch. (**H**) Rat stomach 4 weeks after sutureless repair by the GI patch. (**I** and **J**) Representative histological images stained with hematoxylin and eosin (HE) (left) and immunofluorescence images (right) for rat stomach defect repaired by sutures (I) and the GI patch (J) after 4 weeks. * represents the implanted sutures (D and I) and GI patch (E and J). In immunofluorescence images, blue fluorescence corresponds to cell nuclei stained with 4′,6-diamidino-2-phenylindole (DAPI); green fluorescence corresponds to the expression on fibroblast (αSMA) and macrophages (CD68); red fluorescence corresponds to the expression on collagen (Collagen I) and T-cell (CD3); dotted line represents the edge of the GI patch. (**K**) Normalized fluorescence intensity from the immunofluorescence images for αSMA, Collagen I, CD3, and CD68 4 weeks after repair of rat colon (left) and stomach (right) defects by sutures and the GI patch. (**L**) Complete blood count (CBC) of control (healthy animal without surgery) and 4 weeks after repair of rat colon and stomach defects by sutures and the GI patch. WBC, white blood cell; LYMPH, lymphocyte; RBC, red blood cell; Hb, hemoglobin; HCT, hematocrit; PLT, platelet. (**M**) Blood chemistry of control (healthy animal without surgery) and 4 weeks after repair of rat colon and stomach defects by sutures and the GI patch. AST, aspartate transaminase; ALT, alanine transaminase; ALB, albumin; TP, total protein; BUN, blood urea nitrogen; CHOL, cholesterol; GLU, glucose. Values in K-M represent the means ± SD (*n* = 4 in K; *n* = 6 in L and M). *P* values are determined by the Student’s *t* test for the comparisons between two groups and by one-way ANOVA followed by the Tukey’s multiple comparison test for the comparison between multiple groups; ns, not significant; ** *p* ≤ 0.01; *** *p* ≤ 0.001. Scale bars, 10 mm (B,C,G,H); 200 μm (D,E,I,J).

The histological assessment by a blinded pathologist indicates that the defect repaired by the GI patch is healed with a minimal fibrotic cyst formed only on the top of the GI patch (Fig. 4E and J, and fig. S10C and D), while the sutured defect is healed with mild inflammation and fibrosis around the sutures (Fig. 4D and I, and fig. S10 A and B). Notably, other organs (kidney, liver, spleen, lung) in both groups are normal without any sign of inflammation or damage caused by leaks from the GI defects (fig. S11). The immunofluorescent staining (Fig. 4D,E,I,J) and the normalized immunofluorescence intensity analysis (Fig. 4K) for fibroblasts (αSMA), Collagen I, macrophages (CD68), and T-cells (CD3) indicate that the GI defect repaired by the GI patch induced αSMA, CD68, and CD3 expression is comparable to the standard care by sutures while the expression of Collagen I is significantly higher in the sutured repair. This elevated level of Collagen I expression in the sutured repair agrees with the higher degree of fibrosis observed in the histological evaluation compared to the GI patch-based repair (Fig. 4D,E,I,J and fig. S10). The blood analysis based on CBC (Fig. 4L) and blood chemistry (Fig. 4M) after 4 weeks post-repair does not show significant difference between the health rat control group and the GI defect-repair groups, further confirming that both sutures and the GI patch can prevent anastomotic leaks and subsequent systemic inflammation during the healing of the GI defects (*26*).

### Sutureless repair and healing of GI defects in porcine models

To further validate the *in vivo* efficacy of the GI patch in a more clinically-relevant setting, we demonstrate sealing and healing of two adjacent 5-mm diameter porcine colonic defects per pig by the GI patch (Fig. 5A and B). A total of five pigs received a combined ten 5-mm diameter colon injuries. Notably, we test the GI patch both with and without removable liner based on the *in vivo* porcine colonic defect-repair model (fig. S1B), because it is critical to choose a user-friendly and clinically efficacious packaging method for the GI patch in the pre-clinical evaluation (Fig. 5C).

**Fig. 5.**
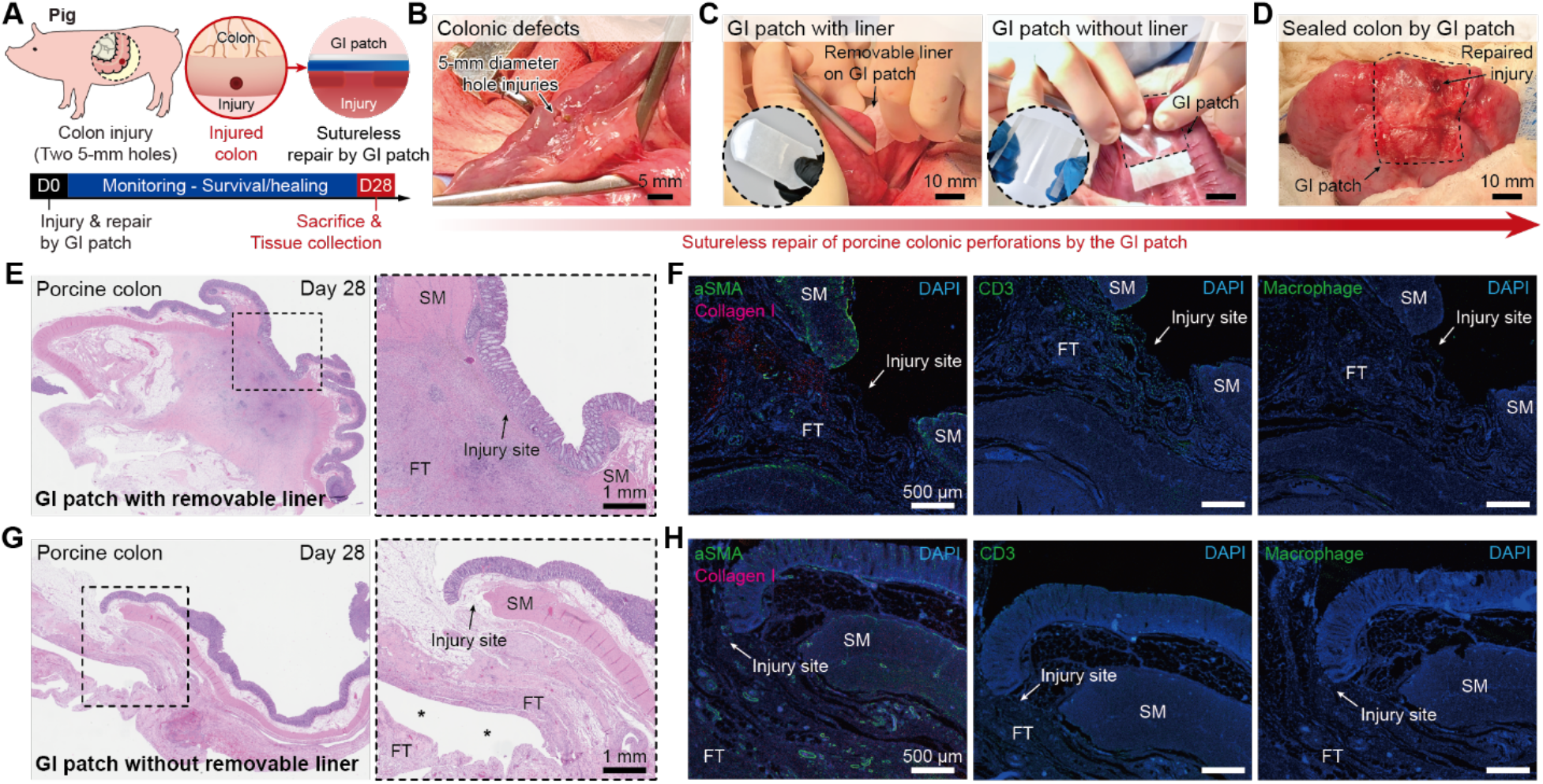
Sutureless repair of GI defects in porcine model. (**A**) Schematic illustrations for *in vivo* sutureless repair of porcine colon defects by the GI patch. (**B-D**) Images for generation of two 5-mm diameter defects in a porcine colon by a biopsy punch (B), sutureless repair of defects by the GI patch with (left) and without (right) removable liner (C), and sealed porcine colon defects by the GI patch (D). (**E** and **F**) Representative histological images stained with hematoxylin and eosin (HE) (E) and immunofluorescence images (F) for porcine colon defects repaired by the GI patch with removable liner after 4 weeks. (**G** and **H**) Representative histological images stained with hematoxylin and eosin (HE) (G) and immunofluorescence images (H) for porcine colon defects repaired by the GI patch without removable liner after 4 weeks. SM, smooth muscle; FT, fibrotic tissue. * represents the GI patch. In immunofluorescence images, blue fluorescence corresponds to cell nuclei stained with 4′,6-diamidino-2-phenylindole (DAPI); green fluorescence corresponds to the expression on fibroblast (αSMA, left), T-cell (CD3, middle), and macrophage (right); red fluorescence corresponds to the expression on collagen (Collagen I). Scale bars, 5 mm (B); 10 mm (C and D); 1 mm (E and G); 500 μm (F and H).

Both GI patch with and without removable liner can provide fluid-tight sealing of porcine colonic defects in less than 10 s (Fig. 5D and movie S5), offering rapid and robust sutureless repair of GI defects. Notably, a relatively steep learning curve is identified by the operating surgeons to achieve full circumferential application of the GI patch around the porcine colonic defects in all cases compared to the rat study. Furthermore, the GI patch without removable liner exhibits a more favorable haptic feedback for the operating surgeons in terms of conformal adhesion to GI defects, owing to the unblocked optical transparency and the lower bending stiffness of the package compared to the GI patch with removable liner.

After repair of a total of 10 potentially lethal colonic defects (two adjacent 5-mm punch holes per pig) by the GI patch, all pigs survived, displayed normal feeding behavior with associated gained weight. There were no signs of abnormal health conditions (e.g., fever or lethargy) or complications associated with wound healing based on daily veterinarian monitoring during the study period of 4 weeks. At the moment of sacrifice after 4 weeks post-repair, no macroscopic signs of GI leaks are observed in any of the animals, despite partial (*n* = 3 animals; repaired by the GI patch without removable liner) or complete detachment (*n* = 2 animals; repaired by the GI patch with removable liner) detachment of the GI patch. It appears that the presence of the removable liner somewhat limited the haptic feedback during the patch application process and may have affected its adhesion performance. The histological evaluation of the repaired porcine colon by a blinded pathologist after 4 weeks indicates that the GI defects are fully healed with fibrotic tissue around them without signs of granuloma or intramural abscess formation for both GI patch with and without removable liner (Fig 5E and G, and fig. S12).

Furthermore, other abdominal organs (liver, spleen, kidney, untreated colon) in both groups are healthy without any sign of inflammation or foreign body reaction (fig. S13). Notably, the defect repaired by the GI patch with removable liner exhibits more fibrosis around the defect in the histological evaluation (Fig. 5E and fig. S12A) and elevated expression of Collagen I, T-Cells (CD3), and macrophages in the immunofluorescent staining analysis (Fig. 5F) compared to the repair by the GI patch without a removable liner (Fig. 5G and H). These findings suggest that both GI patches, with and without removable liner, can provide leak-free sutureless repair and healing of porcine colonic defects. Interestingly, the GI patch without a removable liner appears to yield a more durable attachment and histologically more favorable outcome, which may be due to easier handling and better haptic feedback during the patch application.

## DISCUSSION

While diverse factors contribute to postoperative GI leaks, the mechanical failure of surgically repaired GI defects and subsequent leakage of bowel contents is the most common etiology of GI leaks (*2, 3, 9*). Hence, various treatment strategies have been explored with the aim of improving mechanical sealing of GI defects through tissue adhesive sealants and/or structural reinforcement. For tissue adhesives and sealants, both biologic (e.g., fibrin, gelatin) and synthetic (e.g., PEG, polyurethane, cyanoacrylate) ones have been developed and investigated for repair of GI defects in academic and commercial settings (*27–29*). However, existing tissue adhesives and sealants are fraught with several limitations including mechanical mismatch with GI tissues (e.g., cyanoacrylate), rapid degradation (e.g., fibrin, gelatin), and/or weak sealing strength (*11, 29–32*). Mechanical reinforcement strategies have been explored based on biologic patch (e.g., collagen) or synthetic buttress (e.g., Seamguard, Gore). However, these mechanical reinforcement approaches have shown limited efficacy in pre-clinical studies (*33–35*). Overall, the limitations of existing technologies highlight the unmet clinical needs and the importance of developing new treatment solutions for repair of GI defects.

Mechanical failure of a structure and resultant leakage of fluidic contents within it is also a common problem in non-clinical applications (e.g., pipe leaks). Notably, commercially-available off-the-shelf products such as duct-tapes are frequently used to form almost instant and robust fluid-tight sealing to prevent further leaks, providing both adhesive sealing and mechanical reinforcement to the structural defects (*36*). Despite not yet being developed for and incompatible with clinical and biomedical uses, the remarkable convenience and effectiveness of these off-the-shelf products such as duct-tapes can provide valuable inspiration for the development of novel solutions for the surgical repair of GI defects (*32*).

In this work, we develop an off-the-shelf bioadhesive platform for atraumatic, facile, and robust repair of GI defects to offer a promising solution for unmet clinical needs by addressing multiple limitations of the existing technologies. Inspired by duct-tapes, the GI patch is prepared by synergistically integrating a non-adhesive top layer to provide mechanical reinforcement and a dry bioadhesive bottom layer to offer rapid and robust adhesion to the underlying GI tissue. This unique design along with incorporation of the dry-crosslinking mechanism enables the off-the-shelf, preparation-free and ready-to-use characteristics to the GI patch. We perform systematic characterizations of tissue-matching mechanical properties and adhesion performance of the GI patch based on standard tests and *ex vivo* porcine models. Comprehensive *in vitro* and *in vivo* biocompatibility evaluations show that the GI patch exhibits biocompatibility comparable to that of US Food and Drug Administration (FDA) approved commercially-available tissue adhesive such as Coseal, while being biodegradable long-term (e.g., 12 weeks post-implantation).

Overall, the *in vivo* preclinical rodent and porcine models validate the efficacy of the GI patch as a promising therapeutic device in terms of achieving facile fluid-tight sutureless repair, prevention of leaks, and ultimate healing of the GI defects. However, our study also reveals several limitations and remaining future work to be done. While the GI patch provides facile sutureless sealing of 10-mm incisional intestinal defects in rat models and a total of ten 5-mm diameter colonic punch hole defects in a porcine model, the GI patch would require further validation and optimization for more geometrically and anatomically complex defects. The adhesion performance, packaging, and application procedure of the GI patch may also benefit from further evaluation and optimization in close interaction with practicing surgeons to provide user-friendly haptic feedback and reproducible treatment efficacy. From a biomaterials perspective, the varying degree of inflammation and fibrosis as well as long-term stability of adhesion in different species (i.e., rodent and porcine models) around the GI defects repaired by the GI patch would require future in-depth investigations prior to clinical translation of the technology. Moreover, while the current study supports the *in vivo* biodegradability of the GI patch long-term (e.g., 12 weeks post-implantation), further fine-tuning of the degradation profile while considering the wound healing dynamic of GI defects may offer improved treatment efficacy in the future (*26*). Despite its early developmental stage, the GI patch already offers a promising off-the-shelf platform for atraumatic sutureless repair of GI defects that addresses various limitations of the previous approaches. We envision that the GI patch may not only contribute toward a successful treatment of GI defects but also provide clinical opportunity for repair of other organs and injuries in the human body.

## MATERIALS AND METHODS

### Study design

The aim of this study was to develop an off-the-shelf bioadhesive in the form of ready-to-use, thin, flexible, and transparent patch capable of providing rapid, fluid-tight robust sealing of GI defects with straightforward simple preparation and application. It was hypothesized that a tissue-like bioadhesive material capable of the above-mentioned features can provide atraumatic and effective sealing and repair of GI defects and would address limitations and challenges of surgical repair made by sutures. Systematic mechanical characterizations were performed based on *ex vivo* porcine colon and stomach to evaluate the GI patch’s rapid and robust adhesion to GI tissues with superior adhesion performance in terms of interface toughness, shear strength, tensile strength, and burst strength in comparison with various commercially-available tissue adhesives. *In vitro* LIVE/DEAD assay of human intestinal epithelial cell line (Caco-2) co-culture for 24-h was performed to evaluate cytotoxicity of the GI patch. *In vivo* biocompatibility of the GI patch was assessed based on rat colon and stomach implantation models for 4 weeks followed by histopathological evaluation by a blinded pathologist, immunofluorescence analysis, and blood analysis. *In vivo* biodegradability of the GI patch was investigated based on rat subcutaneous implantation model up to 12 weeks followed by histological evaluations. *In vivo* efficacy of sutureless repair of GI defects by the GI patch was validated by rat colon and stomach defect-repair models and porcine colon defect-repair model for 4 weeks in comparison with a standard care control group based on sutures. The presence of leakage, sealing and healing of GI defects, and overall health of animals were assessed based on animal monitoring, histopathological evaluation by a blinded pathologist, immunofluorescence analysis, and blood analysis.

### Materials

All chemicals were obtained from Sigma-Aldrich unless otherwise mentioned and used without further purification. For preparation of bioadhesive layer of the GI patch, acrylic acid (AAc), poly(vinyl alcohol) (PVA; Mw = 146,000-186,000, 99+ % hydrolyzed), poly(ethylene glycol methacrylate) (PEGDMA; Mn = 550), acrylic acid *N*-hydroxysuccinimide ester (AAc-NHS ester), α-ketoglutaric acid were used. For preparation of the non-adhesive layer of the GI patch, biomedical-grade polyurethane (PU; HydroMed, AdvanSource Biomaterials) was used. As a removable liner for the GI patch, weighing paper (VWR) was used after autoclaving for sterilization. All porcine tissues and organs for *ex vivo* experiments were purchased from a research-grade porcine tissue vendor (Sierra Medical Inc.).

### Preparation of the GI patch

To prepare the bioadhesive layer, 35 w/w % AAc, 7 w/w % PVA, 0.2 w/w % α-ketoglutaric acid, and 0.05 w/w % PEGDMA were added into nitrogen-purged deionized water. Then, 30 mg of AAc-NHS ester was dissolved per 1 mL of the above stock solution to prepare a precursor solution. The precursor solution was then poured on a glass mold with spacers (150 μm thickness for rat study; 350 μm thickness for porcine study) and cured in a UV chamber (354 nm, 12 W power) for 30 min. To introduce the non-adhesive layer, 10 w/w % PU in ethanol solution (ethanol:water = 95:5 v/v) was spin-coated on the as-prepared bioadhesive layer (200 rpm for rat study; 100 rpm for porcine study). To cancel swelling of the GI patch, the as-prepared bioadhesive layer with the spin-coated PU solution was pre-stretched 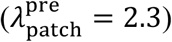 in both length and width directions. The pre-stretched sample was then dried under air flow for 1 h followed by further drying in a vacuum desiccator chamber for 12h to prepare the dry GI patch. A removable liner was introduced for easier handling for rat studies and part of porcine study. The dry GI patch was sealed in a sterile air-tight bag with desiccant (silica gel packets) and stored at −20 °C before use.

### Mechanical characterization

The GI patch was applied to *ex vivo* porcine colon or stomach at 1 kPa pressure (applied either by mechanical testing machine or equivalent weight) for 5 s. All mechanical tests were performed 6 h after initial application of the GI patch to ensure equilibrium swelling of the GI patch. The application of commercially-available tissue adhesives followed the provided user guide or manual for each product. Interfacial toughness was measured based on the standard 180-degree peel test (ASTM F2256, fig. S6A). Shear strength was measured based on the standard lap-shear test (ASTM F2255, fig. S6B). Tensile strength was measured based on the standard tensile test (ASTM F2258, fig. S6C). All tests were performed using a mechanical testing machine (2.5 kN load-cell, Zwick/Roell Z2.5) at a constant crosshead speed of 50 mm min^-1^. Poly(methyl methacrylate) films were applied using cyanoacrylate glue (Krazy Glue) to act as a stiff backing for the GI patch and porcine tissues. Aluminum fixtures were applied using cyanoacrylate glues to provide grips for tensile tests.

### *Ex vivo* demonstration

All *ex vivo* experiments were reviewed and approved by the Committee on Animal Care at the Massachusetts Institute of Technology. To demonstrate rapid and robust sutureless sealing of GI defects by the GI patch, two 5-mm diameter defects were made by a biopsy punch to *ex vivo* porcine colon and stomach. Then, the GI patch was applied on the defects by gentle pressing for 5 s. After repair by the GI patch, saline (red colored by using a food dye) was injected to the porcine colon or stomach to evaluate fluid-tight sealing of the defects. To measure burst pressure, PBS was injected to the sealed porcine colon at the rate of 2 mL min^-1^ while the applied pressure was monitored by a pressure gauge (Omega) (modified ASTM F2392-04, fig. S6D).

### *In vitro* biocompatibility

To evaluate *in vitro* biocompatibility and cytotoxicity of the GI patch, LIVE/DEAD assay was used to assess human intestinal epithelial cell line (Caco-2, ATCC). To prepare conditioned media, 500 mg of Coseal, Histoacryl, and the swollen GI patch were incubated in 10 mL Dulbecco’s modified eagle medium (DMEM) supplemented with 10 v/v % fetal bovine serum (FBS) and 100 U ml^-1^ penicillin–streptomycin at 37 °C for 24 h. The supplemented DMEM without incubating tissue adhesive was used as a control. Caco-2 cells were plated in confocal dishes (20-mm diameter) at a density of 0.5 × 10^5^ cells (*n* = 4 per each group). The cells were then treated with the control and conditioned media, and incubated at 37 °C for 24 h in 5 % CO_2_ atmosphere. The cell viability was determined by a LIVE/DEAD viability/cytotoxicity kit for mammalian cells (Thermo Fisher Scientific) by adding 4 μM calcein and ethidium homodimer-1 into the culture media. A confocal microscope (SP 8, Leica) was used to image live cells with excitation/emission at 495nm/515nm, and dead cells at 495nm/635nm, respectively. The cell viability was calculated by counting live (green fluorescence) and dead (red fluorescence) cells by using ImageJ (version 2.1.0).

### *In vivo* biocompatibility and biodegradability

All animal studies on rat were approved by the MIT Committee on Animal Care and all surgical procedures and post-operative care were supervised by MIT Division of Comparative Medicine veterinary staff. Female Sprague Dawley rats (225-250 g, Charles River Laboratories) were used for all *in vivo* studies. Before implantation, the GI patch was prepared using aseptic techniques and was further sterilized for 3 h under UV light. Commercially-available tissue adhesives were used as provided in sterile packages following the provided user guide or manual for each product.

For *in vivo* biocompatibility evaluation of the GI patch, the animals were anesthetized using isoflurane (2–3% isoflurane in oxygen) in an anesthetizing chamber prior to the surgery, and anesthesia was maintained using a nose cone throughout the surgery. Abdominal hair was removed and the animals were placed on a heating pad during the surgery. Colon or stomach was exposed via a laparotomy. The GI patch (10 mm in width and 20 mm in length) was applied on the colon or stomach surface by gently pressing by a surgical spatula (*n* = 4). For commercially-available tissue adhesives, 0.5 ml of Coseal (*n* = 4) and Histoacryl (*n* = 4) were injected on the colon or stomach surface. The abdominal wall muscle and skin incision was closed using interrupted sutures and 3-4 mL of warm saline were injected subcutaneously. After 4 weeks following the implantation, 3-5 mL of blood was collected via the cardiac puncture technique per each animal for blood analysis, then the animals were euthanized by CO_2_ inhalation. Colon or stomach tissues of interest were excised and fixed in 10 % formalin for 24 h for histological and immunofluorescence analyses.

For *in vivo* biodegradability evaluation of the GI patch, the animals were anesthetized using isoflurane (2–3% isoflurane in oxygen) in an anesthetizing chamber prior to the surgery, and anesthesia was maintained using a nose cone throughout the surgery. The back hair was removed and the animals were placed over a heating pad for the duration of the surgery. The dorsal subcutaneous space was accessed by a 1-2 cm skin incision per implant in the center of the animal’s back. To create space for implant placement, blunt dissection was performed from the incision towards the animal shoulder blades. The GI patch (10 mm in width and 20 mm in length) was placed in the subcutaneous pocket created above the incision (*n* = 4 for each time point). After 2 weeks, 4 weeks, 8 weeks, and 12 weeks following the implantation, the animals were euthanized by CO_2_ inhalation. Subcutaneous regions of interest were excised and fixed in 10 % formalin for 24 h for histological analyses.

### *In vivo* GI organ defect-repair in rat model

For *in vivo* GI organ defect-repair in the rat model, the animals were fasted for 24 h prior to the surgery to minimize bowel contents in the colon and stomach. The animals were anesthetized using isoflurane (2–3% isoflurane in oxygen) in an anesthetizing chamber prior to the surgery, and anesthesia was maintained using a nose cone throughout the surgery. Abdominal hair was removed and the animals were placed on a heating pad during the surgery. Colon or stomach was exposed via a median laparotomy. The exposed colon or stomach was packed with moistened sterile gauzes before creating a defect to prevent contamination of the abdominal cavity. A 10-mm incisional defect was made to the colon or stomach by using a scalpel and repaired by the GI patch (10 mm in width and 20 mm in length) or sutures (8-0 Prolene, Ethicon) (*n* = 4 for each group). After repair of the defect, warm saline was injected to the colon or stomach by a 32G needle syringe to confirm the fluid-tight sealing. The abdominal wall muscle and skin incision were closed with sutures (4-0 Vicryl, Ethicon). After 4 weeks following the repair, 3-5 mL of blood was collected via the cardiac puncture technique per each animal for blood analysis, then the animals were euthanized by CO_2_ inhalation. Colon or stomach tissues of interest were excised and fixed in 10 % formalin for 24 h for histological and immunofluorescence analyses. All animals in the study were survived and kept normal health conditions based on daily monitoring by the MIT Department of Comparative Medicine (DCM) veterinarian staff.

### *In vivo* colon defect-repair in porcine model

All animal studies on pig were approved by the Mayo Clinic IACUC at Rochester. The animals were fasted for 24 h prior to the surgery to minimize bowel contents in the descending colon. The animals were placed in dorsal recumbency and the abdominal region was clipped and prepared aseptically. A blade was used to incise on the ventral midline and extended using electrocautery when indicated. The linea alba was incised and peritoneum bluntly entered, with the incision extended to match the skin incision. The spiral colon was exteriorized and moist lap sponges were used for isolation. Colonic ingesta was milked away from the intended surgical site, and side biting intestinal clamps applied to isolate a portion of the colonic wall. A 5-mm diameter biopsy punch was used to create two lesions in the colon wall. Then a GI patch was applied and adhered over the biopsied region to create a seal (*n* = 2 animals for the GI patch with removable liner; *n* = 3 animals for the GI patch without removable liner). The colon was thoroughly lavaged and returned to the abdomen, then the entire abdominal cavity was lavaged and suctioned then the celiotomy incision was closed. After 4 weeks following the repair, the animals were humanely euthanized, and the wound region sealed by the GI patch was excised and fixed in 10 % formalin for 24 h for histological and immunofluorescence analyses. All animals in the study were survived and kept in normal health condition based on daily monitoring by the Mayo Clinic Rochester veterinarian staff.

### Histology and immunofluorescence

Fixed tissue samples were placed into 70 % ethanol and submitted for histological processing and hematoxylin and eosin (HE) or Masson’s trichrome (MT) staining at the Hope Babette Tang (1983) Histology Facility in the Koch Institute for Integrative Cancer Research at the Massachusetts Institute of Technology. Histological assessment was performed by a blinded pathologist and representative images of each group were shown in the corresponding figures.

For immunofluorescence, paraffin sections of the fixed tissues were cut at 5 μm thickness and baked in 50 °C overnight. The tissue sections were then deparaffinized, rehydrated with deionized water and underwent antigen retrieval using the Steam method. Then the slides were washed in three changes of PBS-Tween20 for 5 min per each cycle. After washing, the slides were incubated in primary antibodies (1:200 mouse anti-αSMA for fibroblast (ab7817, Abcam); 1:200 mouse anti-CD68 for macrophages (ab201340, Abcam); 1:100 rabbit anti-CD3 for T-cells (ab5690, Abcam); 1:200 rabbit anti-collagen-I for collagen (ab21286, Abcam); 1:40 Mouse anti- CD3 (LifeSpan, LS-C350938); 1:40 Mouse anti-Macrophages (Bio-Rad, MCA2317GA)) diluted with IHC-Tek Antibody Diluent for 1 h at room temperature. The slides were then washed three times with PBS-Tween20 and incubated with Alexa Fluor 488 Donkey Anti-Mouse (1:200, Jackson Immuno Research) or Alexa Fluor 594 Donkey Anti-Rabbit secondary antibody (1:200, Jackson Immuno Research) at room temperature in dark environment for 30 min. The slides were washed in PBS-Tween20 three times for 5 min per each cycle. Then the slides were incubated with fluorescent mounting medium with 4′,6-diamidino-2-phenylindole (DAPI) and sealed the edge with a nail polish. A laser confocal microscope (SP 8, Leica) was used for image acquisition. ImageJ (version 2.1.0) was used to quantify the fluorescence intensity of expressed antibodies. All the images were transformed to the 8-bit binary images, and the fluorescence intensity was calculated with normalized analysis. All analyses were blinded with respect to the experimental conditions.

### Statistical analysis

MATLAB (version R2018b) was used to assess the statistical significance of all comparison studies in this work. Data distribution was assumed to be normal for all parametric tests, but not formally tested. In the statistical analysis for comparison between multiple groups, one-way ANOVA followed by Tukey’s multiple comparison test were conducted with the significance threshold of \**p* ≤ 0.05, \*\**p* ≤ 0.01, and \*\*\**p* ≤ 0.001. In the statistical analysis between two groups, the two-sample Student’s *t* test was used with the significance threshold of \**p* ≤ 0.05, \*\**p* ≤ 0.01, and \*\*\**p* ≤ 0.001.

## Acknowledgments

The authors thank the Koch Institute Swanson Biotechnology Center for technical support, specifically K. Cormier and the Histology Core for the histological processing, and Dr. R. Bronson at Harvard Medical School for the histological analyses.

## Author contributions

H.Y., J.W., C.S.N., and X.Z. conceived the idea for the GI patch. H.Y., J.W., C.S.N., L.G.G., and X.Z. designed the research. H.Y. and J.W. developed the materials and method for the GI patch. H.Y., J.W., and C.S.N. designed the *in vitro* and *ex vivo* experiments. H.Y. and J.W. conducted the *in vitro* and *ex vivo* experiment and analysis. J.W. and H.Y. designed and conducted the *in vivo* rat studies and analysis. C.S.N., T.L.S., and L.G.G. designed and conducted the *in vivo* pig studies and analysis. All authors wrote the manuscript.

## Data and materials availability

All data is available in the main text or the Supplementary Information.

## SUPPLEMENTARY MATERIALS

### Supplementary Figures

**Fig. S1.**
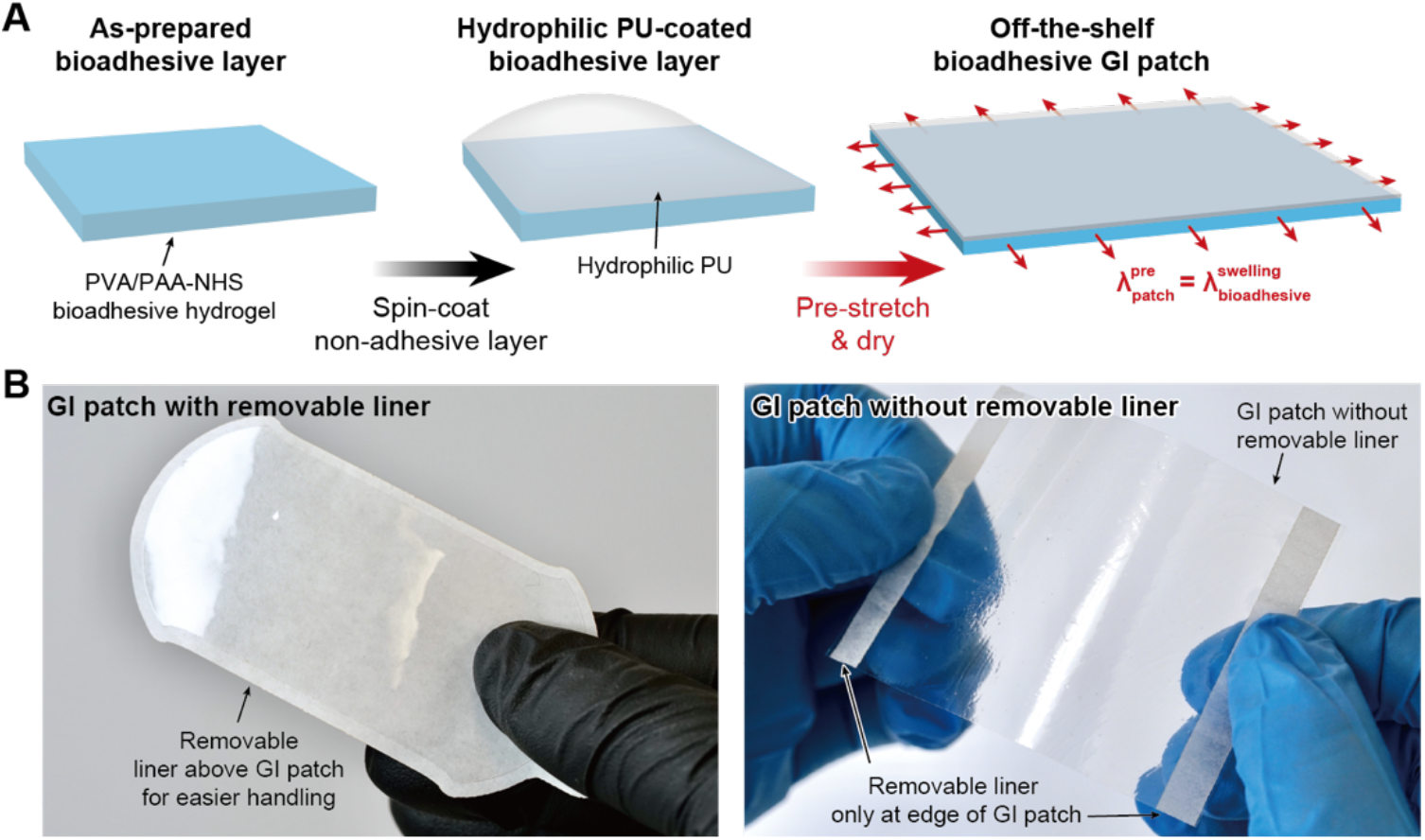
Off-the-shelf bioadhesive GI patch. (**A**) Fabrication process of the GI patch. PU, polyurethane. (**B**) Ready-to-use dry GI patch with (left) and without (right) removable liner.

**Fig. S2.**
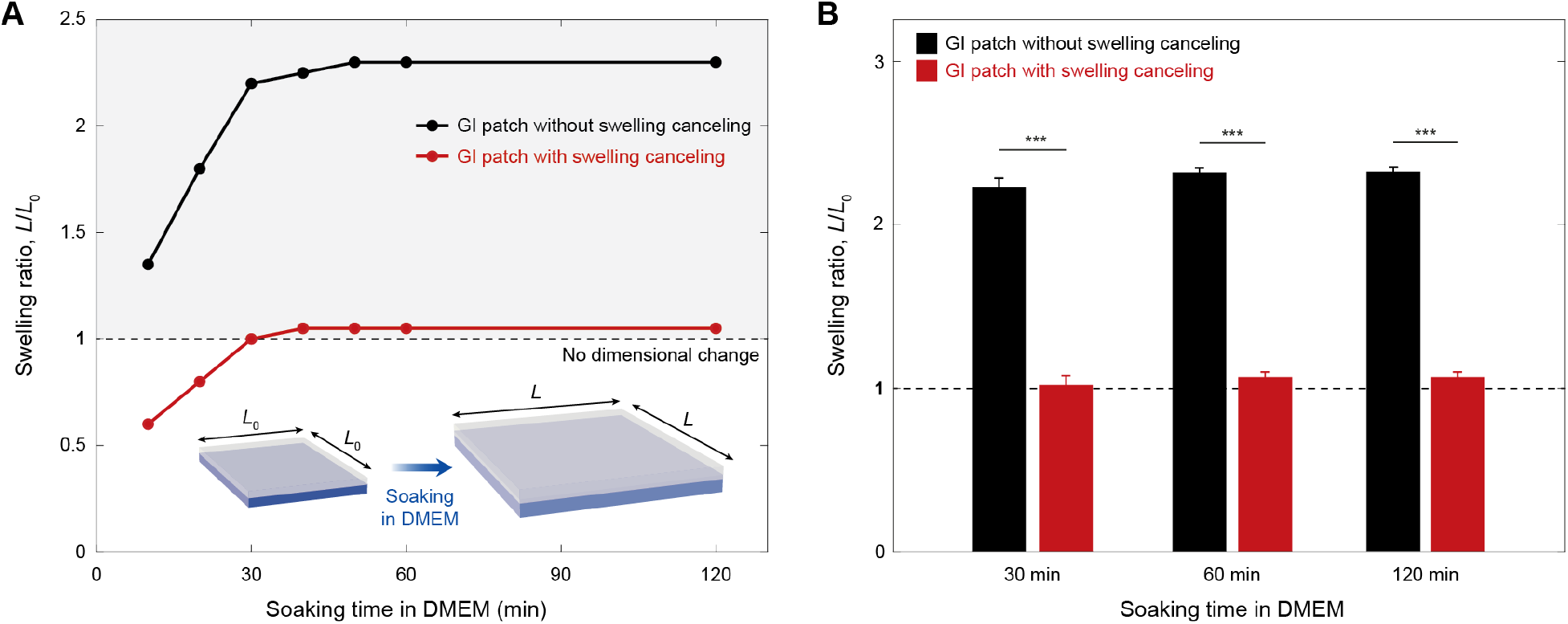
Swelling property of the GI patch. (**A**) Swelling ratio of the GI patch prepared without and with pre-stretch to cancel swelling over time in DMEM at 37 °C. DMEM, Dulbecco’s Modified Eagle Medium. (**B**) Swelling ratio of the GI patch prepared without and with pre-stretch to cancel swelling in DMEM at 37 °C after 30, 60, and 120 min. Values in B represent the means ± SD (*n* = 3). *P* values are determined by the Student’s *t* test; *** *p* ≤ 0.001.

**Fig. S3.**
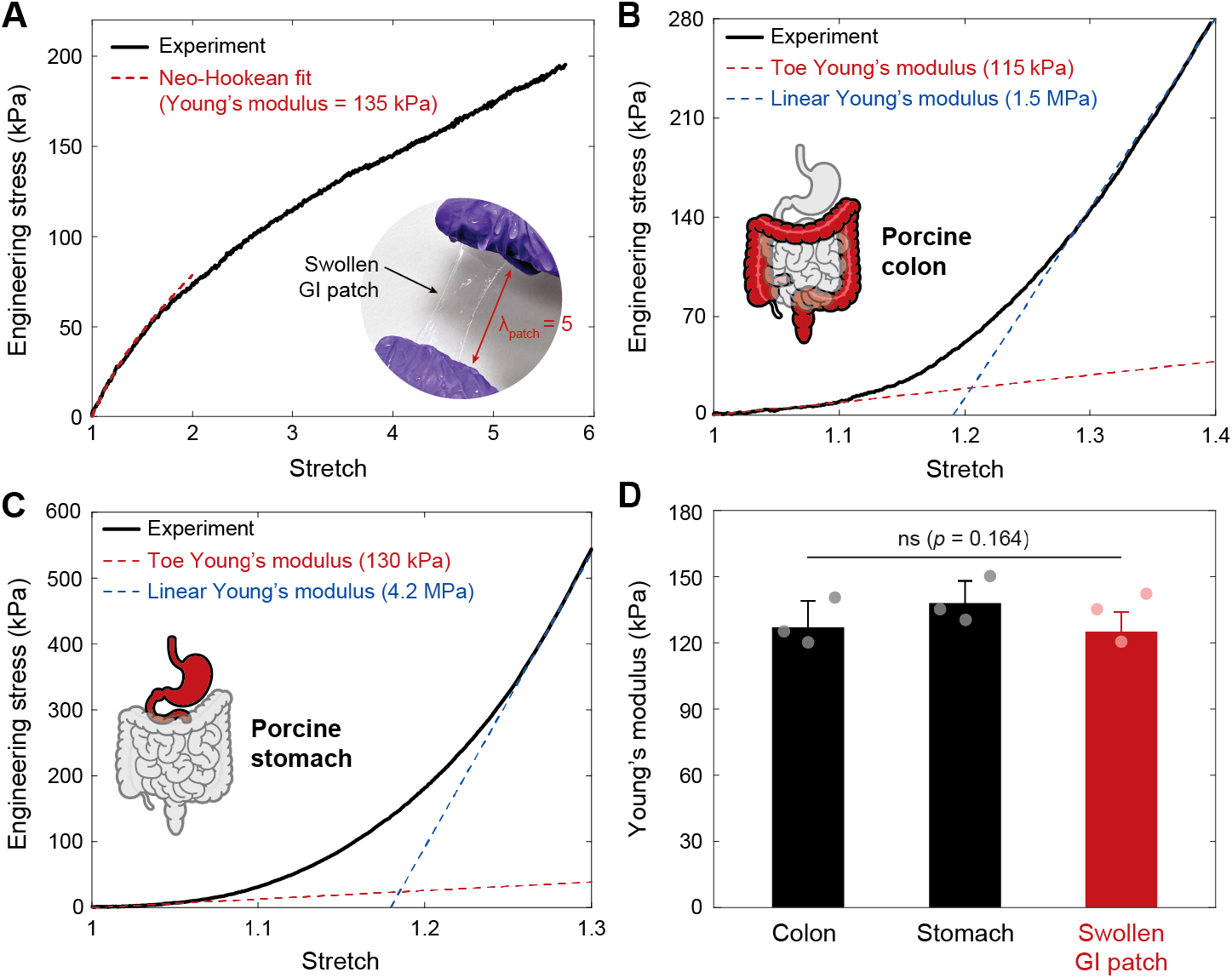
Mechanical properties of the swollen GI patch and porcine GI organs. (**A**) Engineering stress vs stretch curve for the swollen GI patch under tensile deformation. Inset image shows the swollen GI patch stretched 5 times of the original length. (**B** and **C**) Engineering stress vs stretch curves for *ex vivo* porcine colon (B) and stomach (C) under tensile loading. (**D**) Young’s moduli of porcine colon, porcine stomach, and the swollen GI patch. Values in D represent the means ± SD (*n* = 3). *P* values are determined by the Student’s *t* test; ns, not significant.

**Fig. S4.**
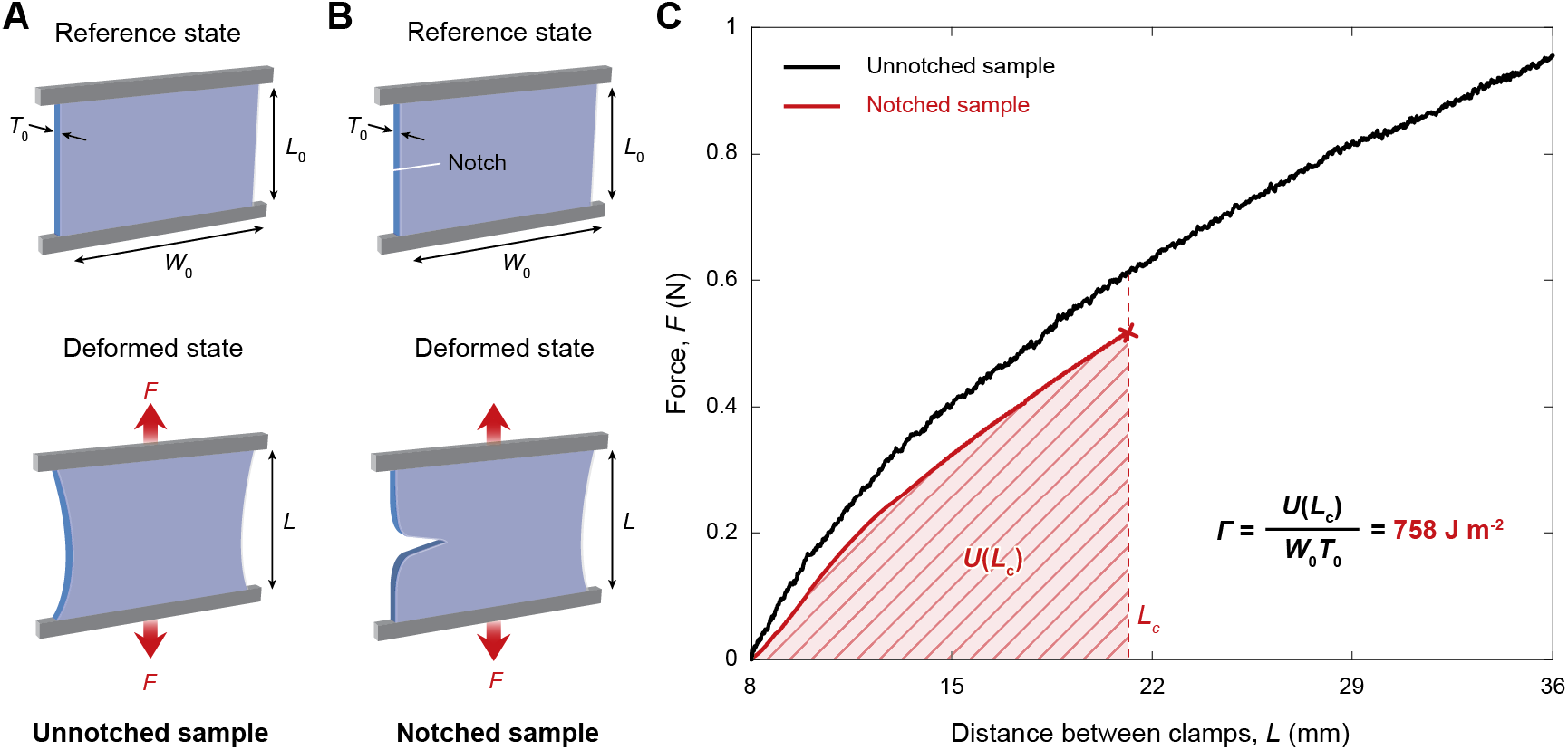
Fracture toughness of the GI patch. (**A** and **B**) Schematic illustrations of pure-shear test for an unnotched sample (A) and a notched sample (B). (**C**) Force vs. distance between clamps for the unnotched and notched swollen GI patch for fracture toughness measurement. *L_c_* indicates the critical distance between the clamps at which the notch turns into a running crack. The measured fracture toughness of the GI patch is 758 J m^-2^.

**Fig. S5.**
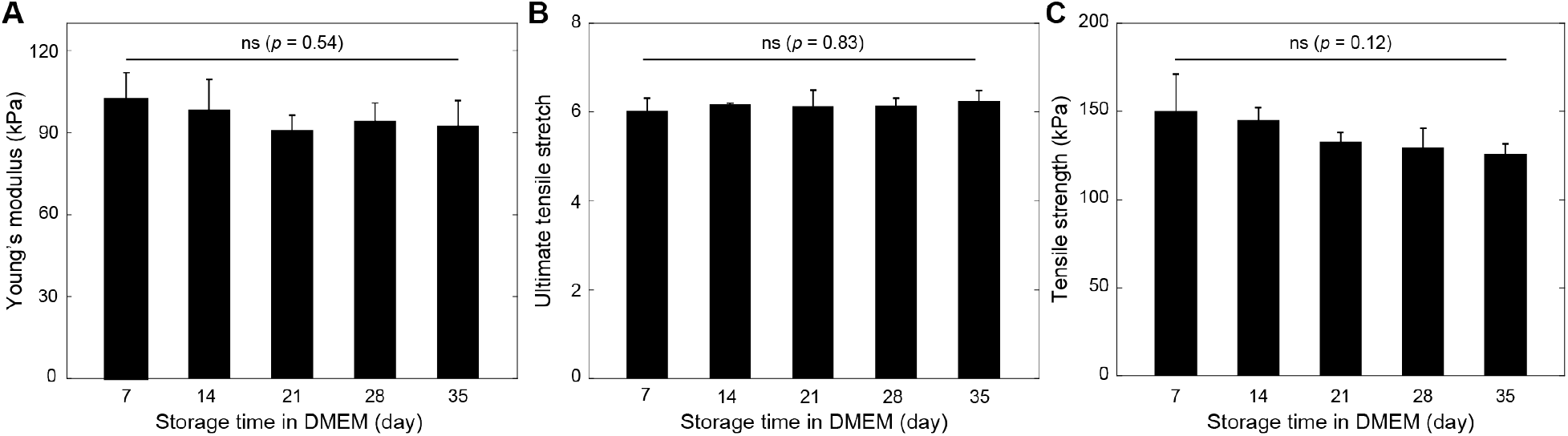
Mechanical stability of the GI patch in physiological environment. (**A-C**) Young’s modulus (A), ultimate tensile stretch (B), and tensile strength (C) of the GI patch stored in DMEM at 37 *°C* over time. DMEM, Dulbecco’s Modified Eagle Medium. Values represent the means ± SD (*n* = 4). *P* values are determined by one-way ANOVA followed by the Tukey’s multiple comparison test; ns, not significant.

**Fig. S6.**
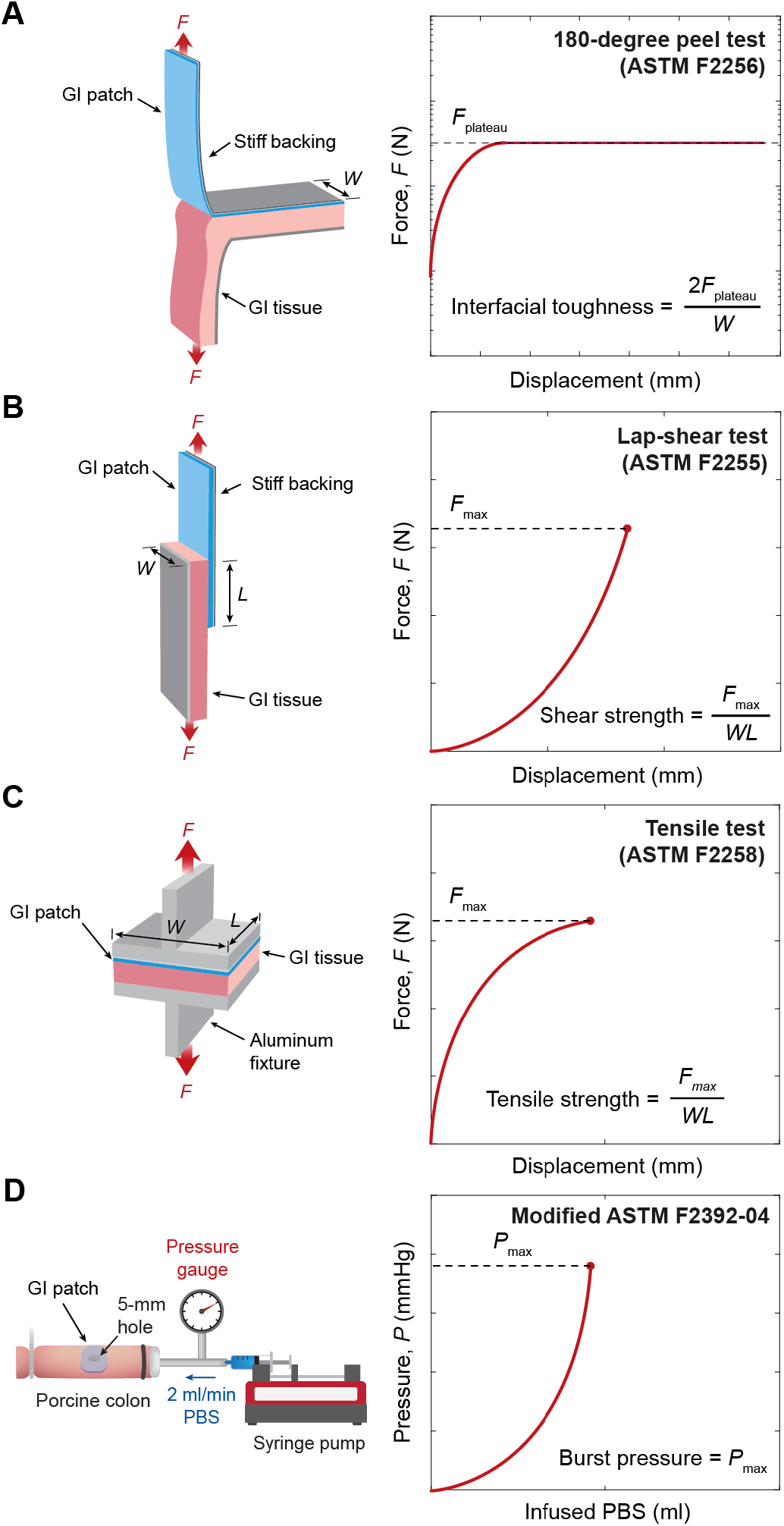
Mechanical testing setups for evaluation of adhesion performance. (**A**) Testing setup for interfacial toughness measurements based on the standard 180-degree peel test (ASTM F2256). (**B**) Testing setup for shear strength measurements based on the standard lap-shear test (ASTM F2255). (**C**) Testing setup for wound closure strength measurements based on the standard tensile test (ASTM F2458-05). (**D**) Testing setup for burst pressure measurements (modified ASTM F2392-04). *F*, force; *W*, width; *L*, length; PBS, phosphate buffered saline.

**Fig. S7.**
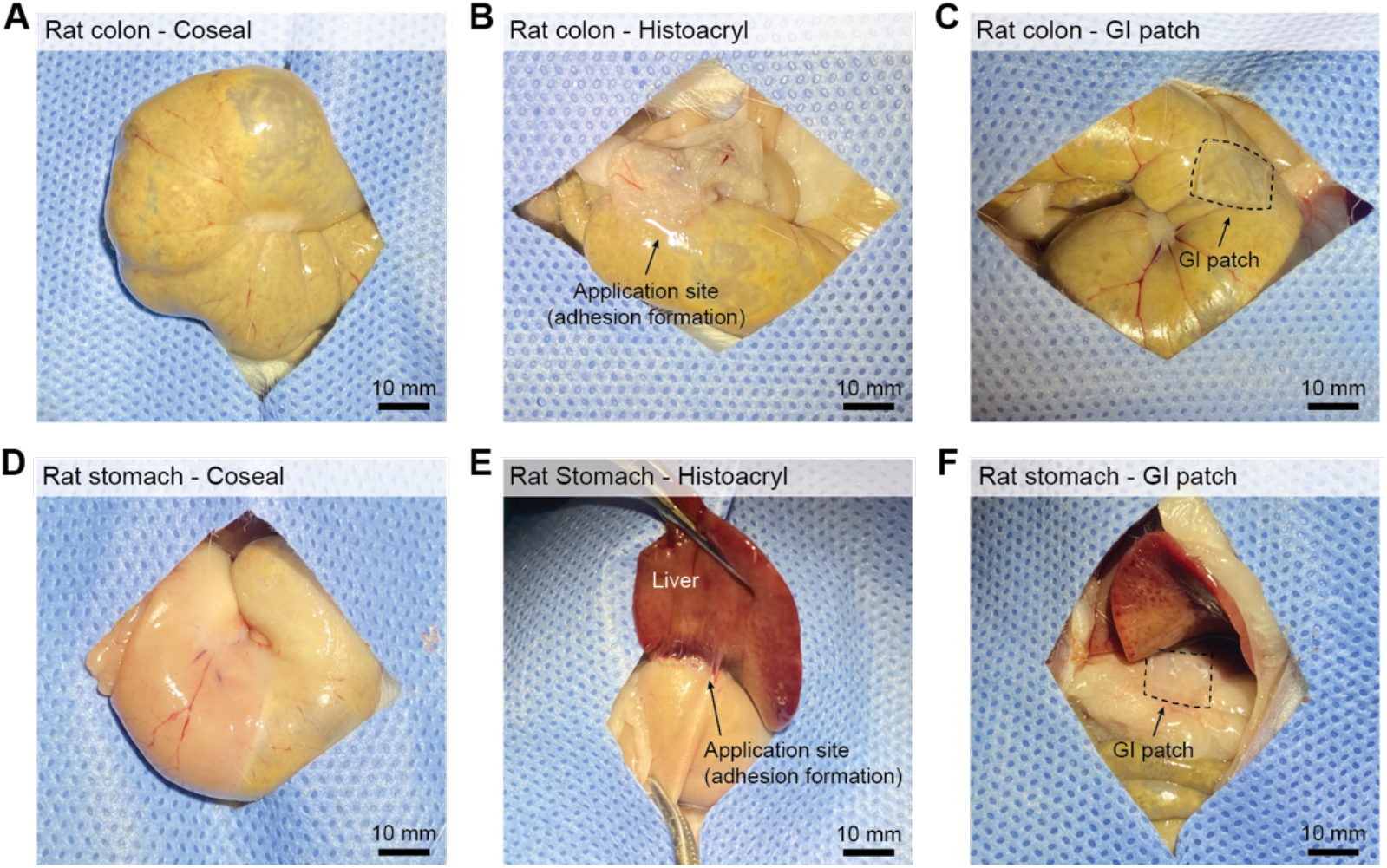
Images of commercially-available tissue adhesives and the GI patch implanted to rat GI organs. (**A-C**) Images of Coseal (A), Histoacryl (B), and the GI patch (C) after 4 weeks implantation to rat colon. (**D-F**) Images of Coseal (D), Histoacryl (E), and the GI patch (F) after 4 weeks implantation to rat stomach. Scale bars, 10 mm.

**Fig. S8.**
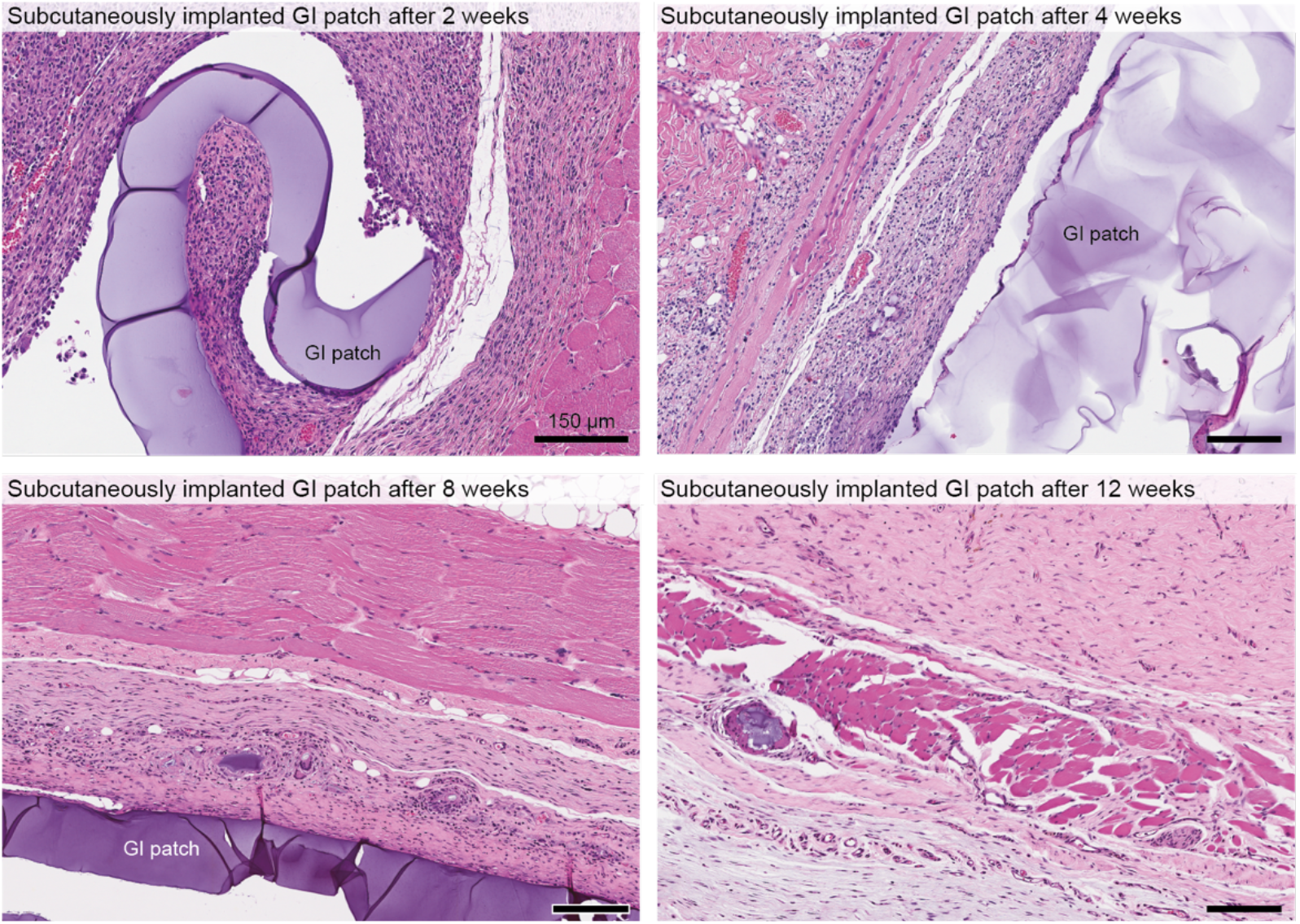
*In vivo* biodegradation of the GI patch in rat model. Representative histological images stained with hematoxylin and eosin (HE) for the GI patch implanted in rat subcutaneous space for 2 weeks (left, top), 4 weeks (right, top), 8 weeks (left, bottom), and 12 weeks (right, bottom). Scale bars, 150 μm.

**Fig. S9.**
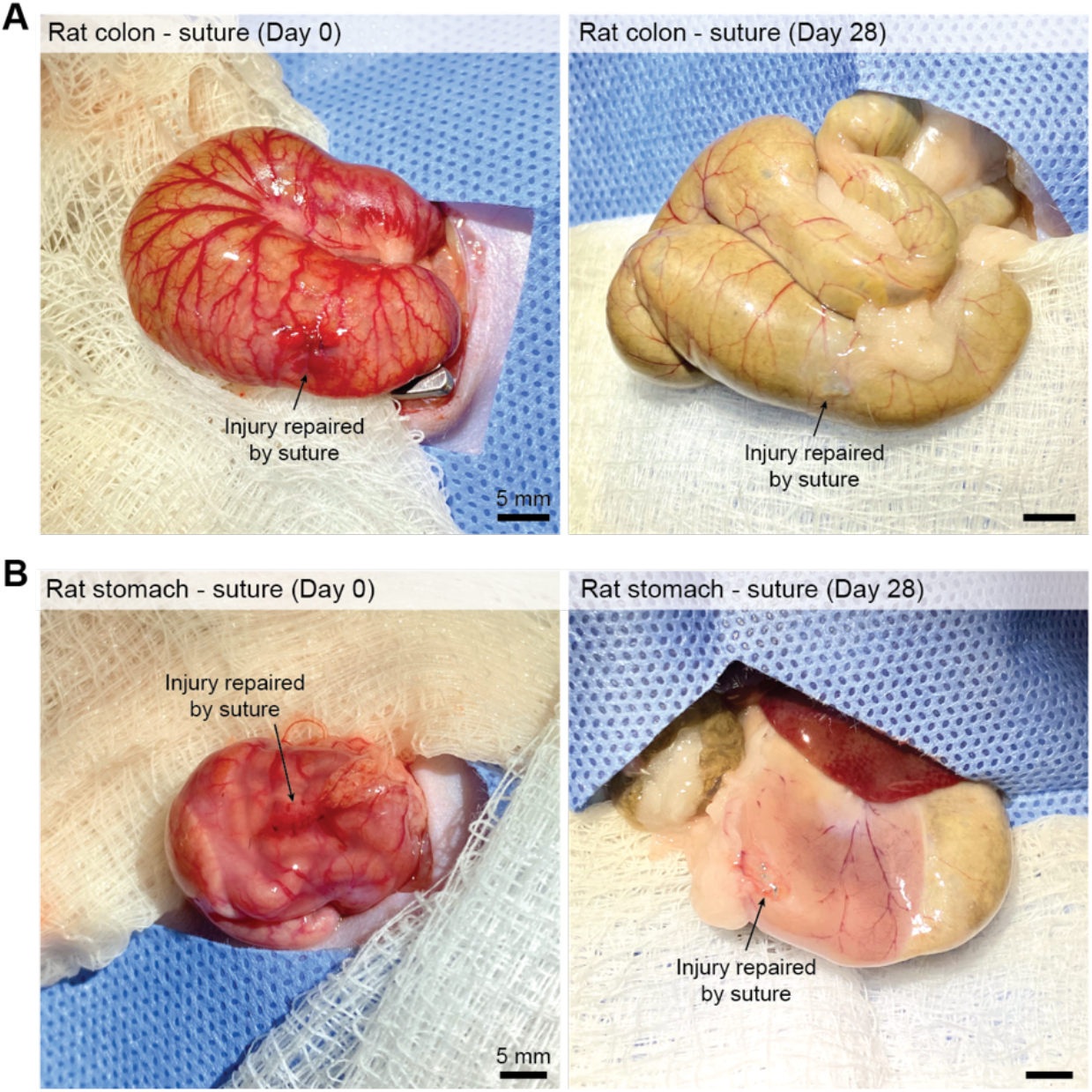
Repair of GI defects by sutures in rat model. (**A** and **B**) Images of rat colon (A) and stomach (B) defects repaired by sutures right after sealing (left) and after 4 weeks (right). Scale bars, 5 mm.

**Fig. S10.**
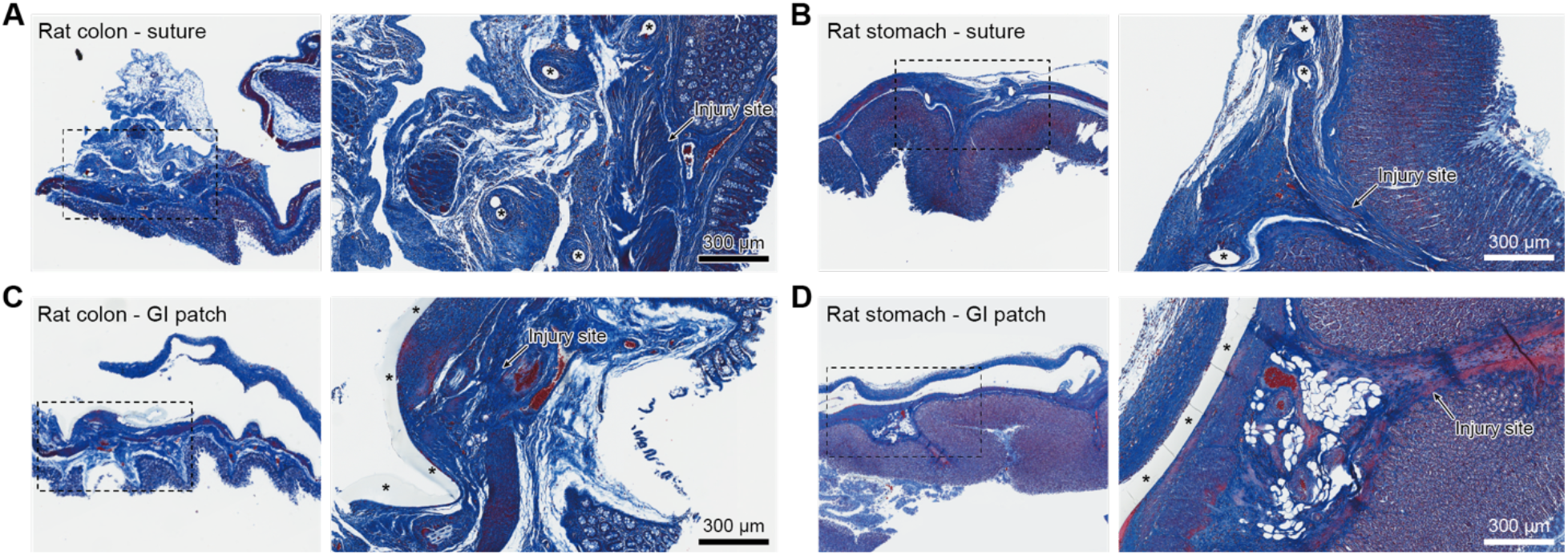
Sutureless repair of GI defects in rat model. (**A** and **B**) Representative histological images stained with Masson’s trichrome (MT) for rat colon (A) and stomach (B) defects repaired by sutures after 4 weeks. (**C** and **D**) Representative histological images stained with Masson’s trichrome (MT) for rat colon (C) and stomach (D) defects repaired by the GI patch after 4 weeks. * represents sutures (A and B) and the GI patch (C and D). Scale bars, 300 μm.

**Fig. S11.**
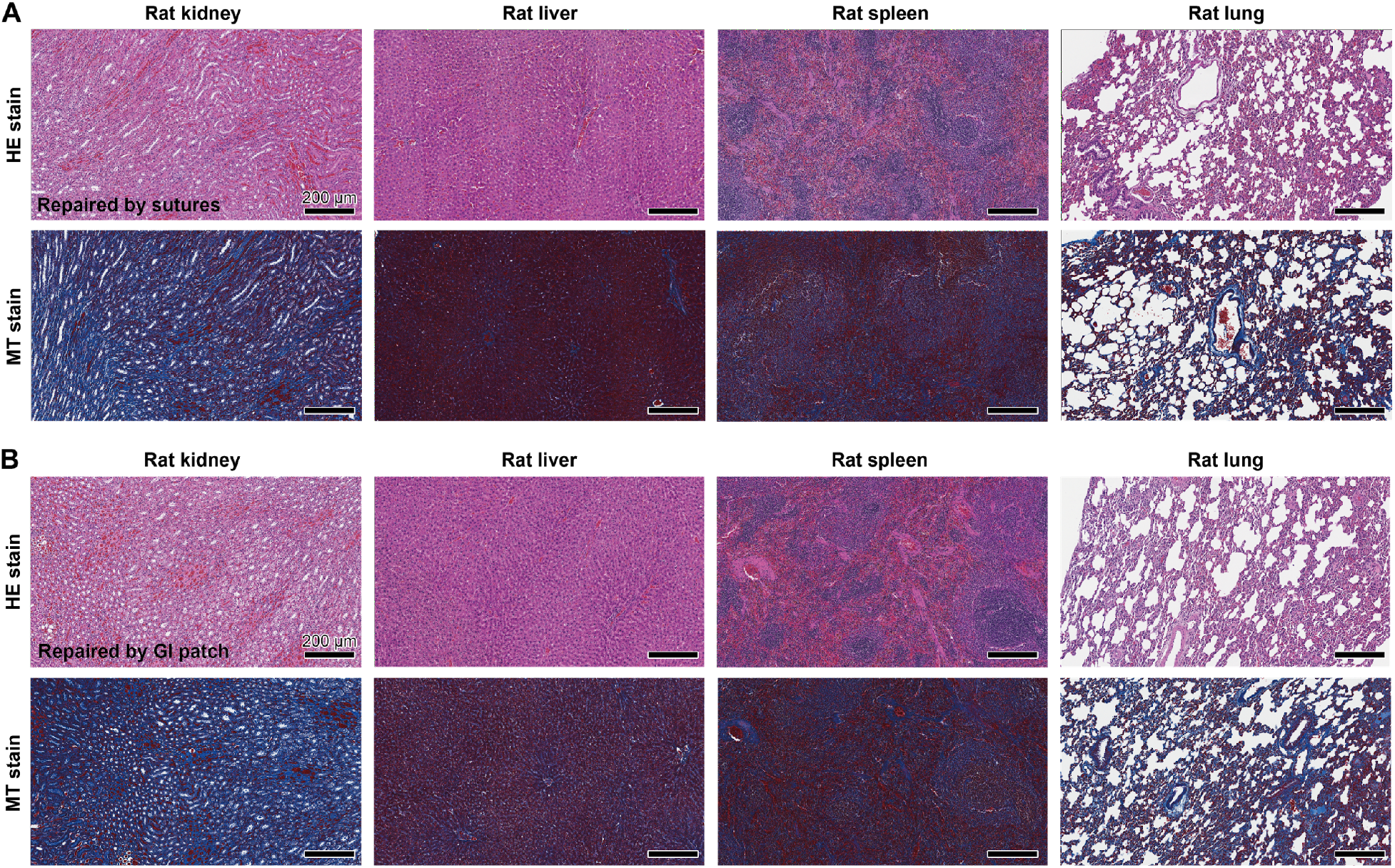
Other organs in rat GI defect-repair study. (**A**) Representative histological images stained with hematoxylin and eosin (HE, top) and Masson’s trichrome (MT, bottom) for other organs in rat with colon defects repaired by sutures after 4 weeks. (**B**) Representative histological images stained with hematoxylin and eosin (HE, top) and Masson’s trichrome (MT, bottom) for other organs in rat with colon defects repaired by the GI patch after 4 weeks. Scale bars, 200 μm.

**Fig. S12.**
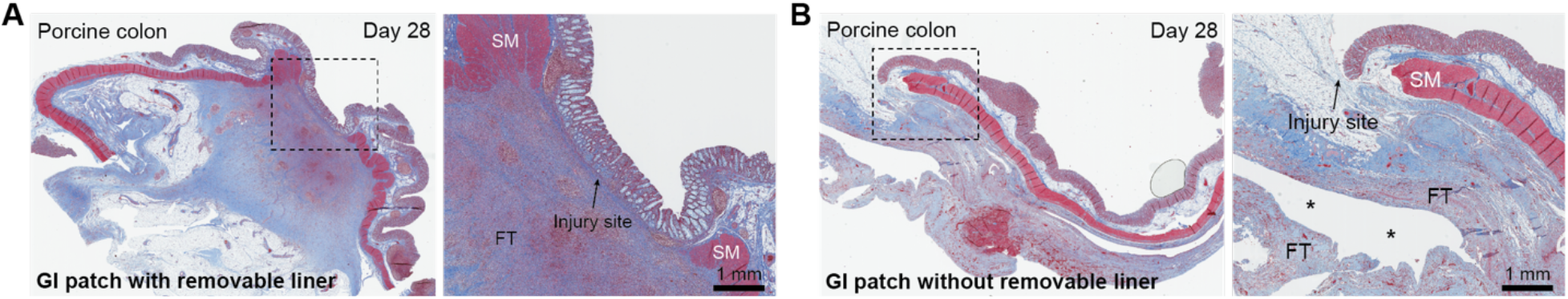
Sutureless repair of GI defects in porcine model. (**A** and **B**) Representative histological images stained with Masson’s trichrome (MT) for porcine colon defects by the GI patch with (A) and without (B) removable liner after 4 weeks. * represents the GI patch. SM, smooth muscle; FT, fibrous tissue. Scale bars, 1 mm.

**Fig. S13.**
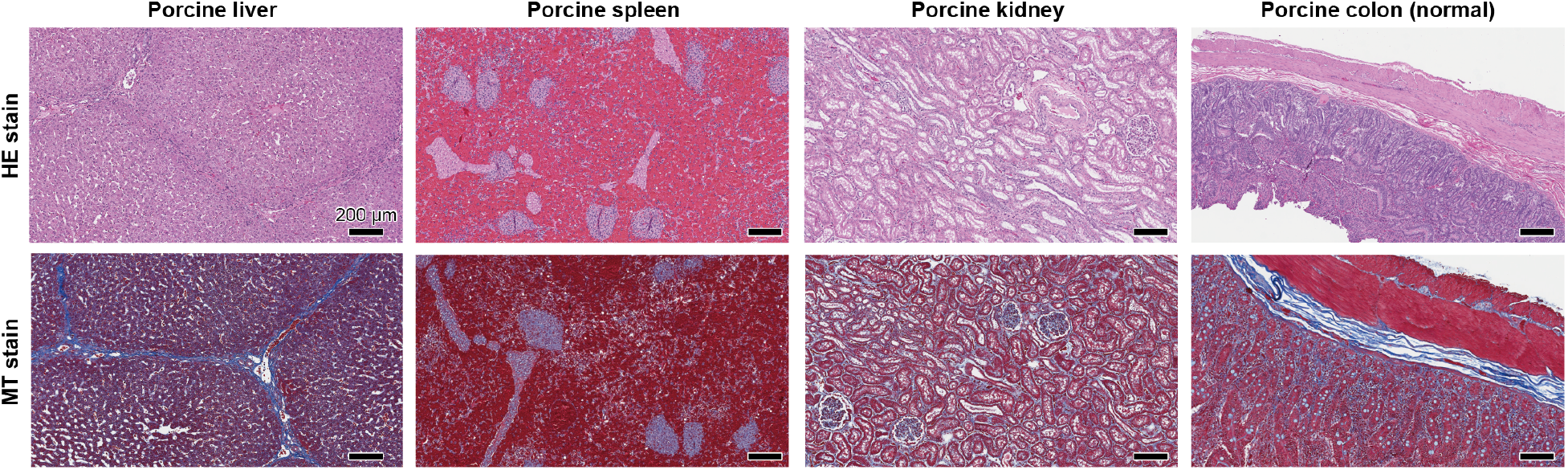
Other organs in porcine GI defect-repair study. Representative histological images stained with hematoxylin and eosin (HE, top) and Masson’s trichrome (MT, bottom) for other organs in pig with colon defects repaired by the GI patch after 4 weeks. Scale bars, 200 μm.

### Captions for other supplementary materials

**Movie S1.** Sutureless repair of a gastrointestinal defect in an *ex vivo* porcine colon by the GI patch.

**Movie S2.** Sutureless repair of a gastrointestinal defect in an *ex vivo* porcine stomach by the GI patch.

**Movie S3.** Sutureless repair of a gastrointestinal defect in an *in vivo* rat colon by the GI patch.

**Movie S4.** Sutureless repair of a gastrointestinal defect in an *in vivo* rat stomach by the GI patch.

**Movie S5.** Sutureless repair of gastrointestinal defects in an *in vivo* porcine colon by the GI patch.

